# The neurovascular unit of capillary blood vessels in the rat nervous system. A rapid-Golgi electron microscopic study

**DOI:** 10.1101/2023.04.26.538454

**Authors:** Jorge Larriva-Sahd, Gema Martínez-Cabrera, Carlos Lozano-Flores, Luis Concha, Alfredo Varela-Echavarría

**Affiliations:** Instituto de Neurobiología Universidad Nacional Autónoma de México, Campus Juriquilla, Querétaro, México

**Keywords:** perivascular synapse, BOLD, Piezo 1, mechanosensory

## Abstract

We describe a pericapillary organ in the rat forebrain and cerebellar cortex. It consists of series of tripartite synapses enveloped by astrocytic endfeet linked to the capillary wall by synaptic extensions. Reciprocal specializations of the pericyte-capillary blood vessel with such specialized synapses suggests a mechanoreceptor role. In Golgi impregnated and 3D reconstructions of cerebral cortex and thalamus, series of tripartite synapses appear sequentially ordered in a tributary dendrite paralleled by synaptic outgrowths termed here golf club-like extensions apposed to a longitudinal crest from the capillary basal lamina. To facilitate identification of principal cell dendrites and arriving axons to these putative mechanosensory structures, we utilized the cerebellar cortex since it has a well known organization and observed that afferent fibers and interneurons display interactions with the capillary wall. Afferent mossy fiber rosettes and ascending granule cell axons and dendrites define pericapillary on passage interactions surrounded by endfeet. The ability of such structures to modulate synaptic transmission is supported by the presence of mRNA of the mechanosensitive channel Piezo 1 in the mossy fiber rosettes, pyramidal isocortical and thalamic neurons. This suggests that ascending impulses to the cerebellar and cortical targets are presynaptically modulated by the reciprocal interaction with the mechanosensory pericapillary organ.

## Introduction

Brain function relies on the adaptation of blood supply to neural and glial activity. Processes involved in the control of the brain circulation are too important to be overlook (Roy and Sherrington, 1890). Because of the viability of the highly differentiated nerve cells relies upon exchange of bioactive molecules with capillary blood vessels this process is of fundamental relevance. While peripheral blood circulation appears to be chiefly controlled by the autonomous nervous and endocrine systems, cerebral blood supply is driven by different regulatory mechanisms (Roy and Sherrington 1890). More specifically, researchers assumed that the motor control of capillary blood flow relies on cranial nerve and central aminergic systems innervating tributary muscular arteries since no meningeal nerve fibers penetrate beyond Robbin-Virchow’s space (Jones, 1970). Hence, the capillary blood flow results from the passive reception of arterial blood while motor activity is virtually absent. Fundamental for the present study is the widespread observation that neuronal or receptor (Chaigneau, et al., 2003) recruitment within discreet areas of the cerebral cortex leads to a rapid increase of blood flow that ceases as the stimulus ends (Cauli and Hamel 2010). This neurovascular coupling is the cornerstone of functional magnetic resonance imaging (refs Grinvald et al., 1991, Bandettini, et al., 1992, Ogawa, et al., 1992). While the mechanisms that initiate the hemodynamic response are documented (refs Atwell, Iadecola 2017, McCaslin, et al., 2011), an important question remains: How does the intended effect (i.e., increase in blood flow) feed back to the motor pool?. Incidentally, most motor and endocrine systems, initially thought to be exclusively efferents have, as a rule, feed-back mechanisms via sensory organs, or hormones, respectively. In keeping with this, it has been assumed that is the orchestrated interaction of endothelial cells, astrocytes and neurons is what enables the subtle capillary blood flow variation, to tune the motor limb of the circuit (Idecola 2004, Zonta et al., 2002; Tsvetanov et al., 2020, Idecola 2017, Moore and Cao 2008,. Idecola and Nedergard 2017, Armulik et al, 2010, Reichenbach and Wolburg, 2009), researchers nonetheless fail defining substrate(s) linking them. More precisely, which sectors of the vascular tree between arterioles and terminal capillaries are responsible for CBf regulation is a matter of debate (Idecola and Nedergard 2007, Armulik et al, 2010, Idecola 2017). Over the past decade we have investigated the perivascular neuropil and described first a set of neurons within the brain vasculature, specifically with arteries. The long processes of these unique perivascular neurons (PVNs) associated with granule-containing, tuberous structures at sites of ramifying blood vessels that appear to correspond to mechanoreceptors (Larriva-Sahd et al., 2019). However, given than the PVN and its processes are absent in capillaries with diameter of 15µm or less (Larriva-Sahd et al., 2019), we decided to study the structure of the brain-capillary blood vessel interphase. While the EM-3D descriptions of capillary blood vessels and associated neural and glial processes are available (Reichenbach, et al. 2010, Reichenbach and Wolburg 2008, Mathiisen, et al 2010, Shapson-Coe, et al 2021) a systematic search for structural interactions between them is necessary. Hence, the first goal of present study was to revisit the structure of the capillary wall and neighboring neuropil with the light and electron microscope. This allowed the identification of a novel series of synaptic, dendritic, and astroglial specializations that reciprocate with the capillary wall, their neuronal sources, and obtained an initial molecular characterization.

## Material and Methods

### Golgi Technique

Brains from normal, pathogen-free adult Wistar rats (n=25) raised in our vivarium facility used as in our previous studies (Larriva-Sahd 2006, 2008), perivascular cells, interstitial ground substance, and nerve fibers structuring the neurovascular unit (Hawkins and Davis 2008) were studied. Briefly, after sacrifice with a barbiturate overdose, brains were removed from the skull and transversely cut into equidistant quarters with a sharp pair of scissors. Tissue blocks approximately 4mm-thick were sampled from the brain stem; cerebellum; frontal and parietal isocortices; hippocampus-entorhinal cortex; diencephalon; basal ganglia; olfactory cortex; olfactory bulb; and peduncle. Each sample was placed in an aqueous solution containing 0.25% osmium tetroxide and 3% potassium dichromate for 10 to 13 days and transferred to a 0.75% silver nitrate solution for incubation 20 additional days. Each block of tissue was encased with a paraffin shell and serially sectioned at a thickness of 150 microns with a sliding microtome. Sections were transferred to 70% ethyl-alcohol, dehydrated, cleared with Terpineol-xylene, mounted, numbered, and coverslipped with Entellan (Merck). Species conservation of the pericyte and its processes (Figs. 10 and 11), was assessed in specimens of adult mouse, rat, hamster, rabbit, and rhesus monkey from our archives. Images were obtained with a camera lucida adapted to an Axioplan 2 (Zeiss, Oberkochen) light microscope, using 40X and 100X oil objectives. Indian ink and soft pencil drawings were obtained from representative areas. Somatic, axonal, dendritic, and vesicular structures were measured in digital images acquired with an AxioCam camera using Kontron software (Zeiss, Oberkochen). Illustrative light microscopic images from Golgi-impregnated specimens were obtained by overlapping a variable number of in sharp focus areas into a single micrograph.

### Electron Microscopy

Brains from five adult rats (vide supra) were used for this part of the study. Subjects were deeply anesthetized and perfused through the left ventricle with Karnovsky’s fixative and processed as previously described (Larriva-Sahd, 2006, 2008 Varela-Echavarría et al., 2017). Tissue blocks approximating 1mm thick from the frontal and parietal isocortices, posterior thalamus and cerebellar cortex were dissected out, according to Swanson’s atlas (Swanson,). Samples of tissue were postfixed for one hour in 1% osmium tetroxide dissolved in 0.1 M cacodylate-HCl buffer; then they were dehydrated in graded acetone and embedded in epoxy resin. One micrometer-thin sections were obtained from the tissue blocks in a Leica ultramicrotome equipped with glass knives. Silver to gold-colored sections were obtained with a Leica ultramicrotome equipped with glass knives and mounted on copper grids. Then, sections were sequentially stained with uranium and lead salts and they were observed under a JOEL 1010 electron microscope.

**Table 1.**
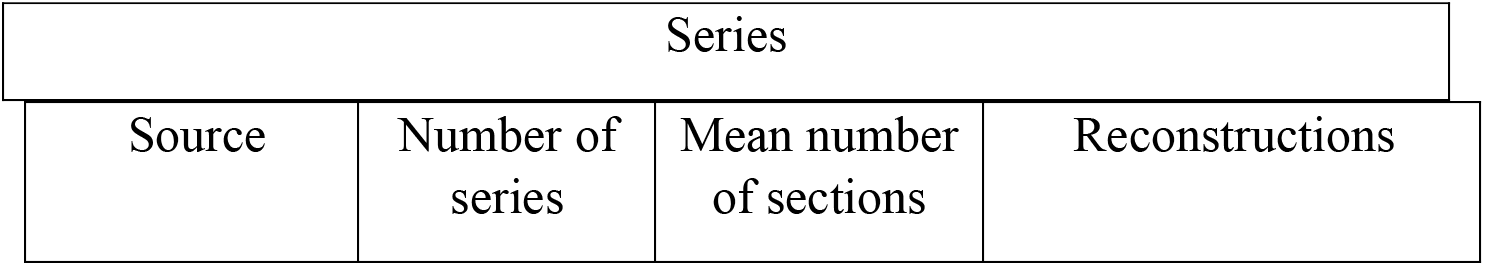
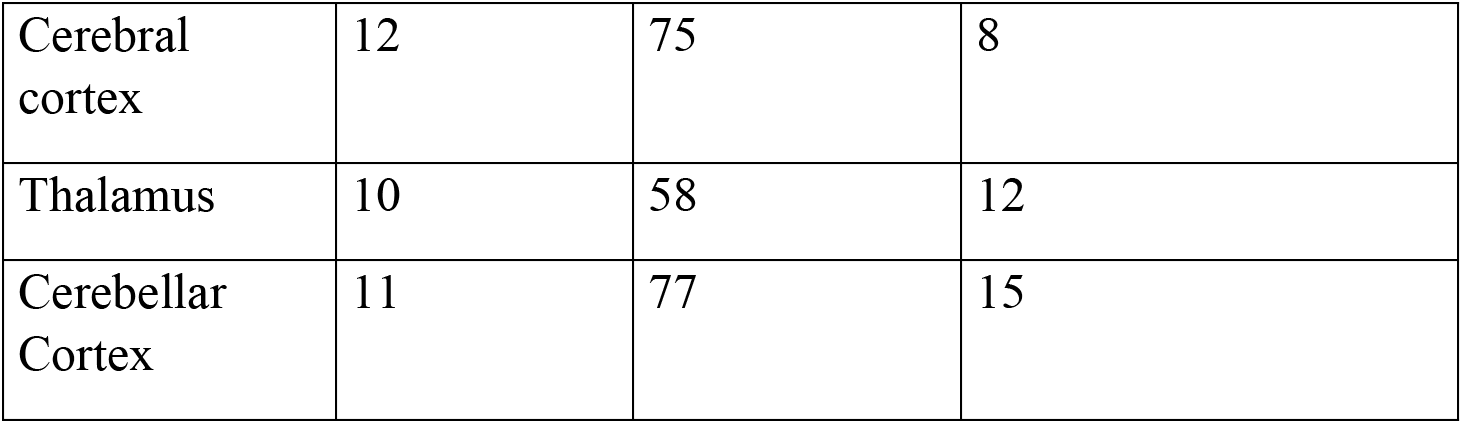
Series observed at the electron microscope and 3D reconstructions.

| Series |  |  |  |
| --- | --- | --- | --- |
| Source | Number of series | Mean number of sections | Reconstructions |
| Cerebral cortex | 12 | 75 | 8 |
| Thalamus | 10 | 58 | 12 |
| Cerebellar Cortex | 11 | 77 | 15 |

Serial Sectioning and 3D Reconstructions. For tridimensional reconstructions tissue blocks from the adult rat hippocampus, cerebral-, cerebellar cortex, primary olfactory cortex, thalamus, and olfactory bulb were used (Table 1). Series were obtained with the Leica microtome equipped with a diamond knife. Successive 70 to 80 nanometers thin sections were mounted on one-slot cupper grids that had previously been covered with Formvar film. Then, specimens were sequentially stained with aqueous uranium and lead salts and studied at the electron microscope. Areas of interest were serially photographed at the electron microscope with a Gatan digital camera. Images from each series were aligned first with the ImageJ-FIJI software, free-hand outlined with a cursor, and then reconstructed with a software that is freely available (Reconstruct, http://synapses.clm.utexas.edu/tools/reconstruct/reconstruct.stm.). We used microtome diamond-cut series as they provide the superior resolution of the transmission EM as well as the invaluable opportunity to perform post hoc observation that otherwise are lost in gallium sectioned specimens which are sublimated and therefore lost following observation.

### Immunohistochemistry and *In situ* Hybridization

In situ hybridization. A fragment of the rat Piezo 1 cDNA was amplified with Taq Polymerase and the oligonucleotide primers 5’-GAGGAAGAGGACTACCTT and 5’-TTTACTTAGAAAACCCTACAG from bladder total RNA and cloned into the pGEM-T Easy vector (Promega, Madison, WI). Sequence was confirmed by Sanger capillary sequencing and sense and antisense RNA probes were synthesized with T7 and SP6 RNA polymerases (Roche-Sigma-Aldrich), respectively and FITC-UTP (with FITC RNA labelling kit from Roche). These probes were then used for in situ hybridization on cryosections of fresh brain tissue which was snap-frozen in blocks of Tissue-Tek (Sakura) employing an isopentane bath cooled with powdered dry ice. Cryosections of 30 microns were obtained on SuperFrost Plus slides (FisherScientific) and fluorescent in situ hybridization (FISH) was performed as described (Hernández-Linares, Y et al 2019 (doi: 10.1038/s41598-019-43701-w). The FITC-UDP detection was using peroxidase conjugated anti-FITC antibody (Roche) and FITC-tyramide signal amplification system (Akoya bioscience).

Fluorescent immunostaining was performed on cryosections obtained on Superfrost Plus slides of tissues fixed in 3.5% paraformaldehyde in PBS employing the antibodies of Tables1 and 2. Immunstaining of vGlut and CGRP was performed in cryosections of cerebellum of mice expressing the reporter GFP protein under the control of the GFAP promoter (Nolte et al., 2001) and DAPI staining for nucleus and IB4 for capillary blood vessels.

The fluorescence was detected using a Confocal Microscope Carl Zeiss – LSM 880 and ZEN software and Plan-Apo 63x N.A.=1.4 oil immersion objetive. The 3D images were obtained in Z-stacks from 30 to 40 µm of depth with 1024 × 1024 pixels of resolution and analyzed with the image-processing package Fiji (version: 2.0.0-rc-66/1.52b; http://imagej.net/contributors) of the Image J program and Amira Software (https://www.thermofisher.com/mx/es/home/electron-microscopy/products/software-em-3d-vis/amira-software.html).

**Table 2.** List and characteristics of primary antibodies to immunohistochemistry.

| Primary Antibodies |  |  |  |  |  |
| --- | --- | --- | --- | --- | --- |
| Name | Host | Clonality | Dilution | Catalog #/ID | Manufacturer |
| VGlut2 | Mouse | Monoclonal | 1:500 | Ab79157/AB1603114 | Abcam |
| Anti-Fluorescent<br>POD | Sheep | Monoclonal | 1:1000 | 11426346910/AB-84057 | Sigma/Roche |

**Table 3.** Secondary antibodies used to visualize immunoreactive sites to primary antibodies.

| Secondary Antibodies |  |  |  |  |  |  |
| --- | --- | --- | --- | --- | --- | --- |
| Name | Host | Conjugate | EX/EM (nm) | Dilution | Catalog #/ ID | Manufacturer |
| Anti-mouse | Donkey | Alexa-Fluor<br>594 | 590/617 | 1:1000 | Ab150112/<br>Ab-2813898 | Abcam |

The fluorescence was detected using a Confocal Microscope Carl Zeiss – LSM 880 and ZEN software and Plan-Apo 63x N.A.=1.4 oil immersion objetive. The 3D images were obtained in Z-stack from 30 to 40 µm of depth with 1024 × 1024 pixels of resolution and analysed with the image-processing package of Fiji (version: 2.0.0-rc-66/1.52b; http://imagej.net/contributors) of the Image J program and Amira Software (https://www.thermofisher.com/mx/es/home/electron-microscopy/products/software-em-3d-vis/amira-software.html).

### Nomenclature and information retrieval

We adopted rigid, accepted criteria to define capillary blood vessels CBVs. At the light microscopy CBV plexuses are considered those arising from terminal arterioles of less than 15 microns (McDonald and Larue, 1983) and leading to postcapillary veins. The higher affinity for silver and smaller caliber of the former, allow unambiguous differentiation from the later (Fig. 1A). At the EM, the CBV is considered to be that vessel having a diameter of <5 micrometers, *and* surrounded by pericytes whose cytoplasm is therefore devoid of myofibrils and/or electron-dense submembrane clusters of myofibrils, typically observed in myocytes (Hill et al., 2015). With these guidelines a consistent CBV identification was performed.

**Figure 1.**
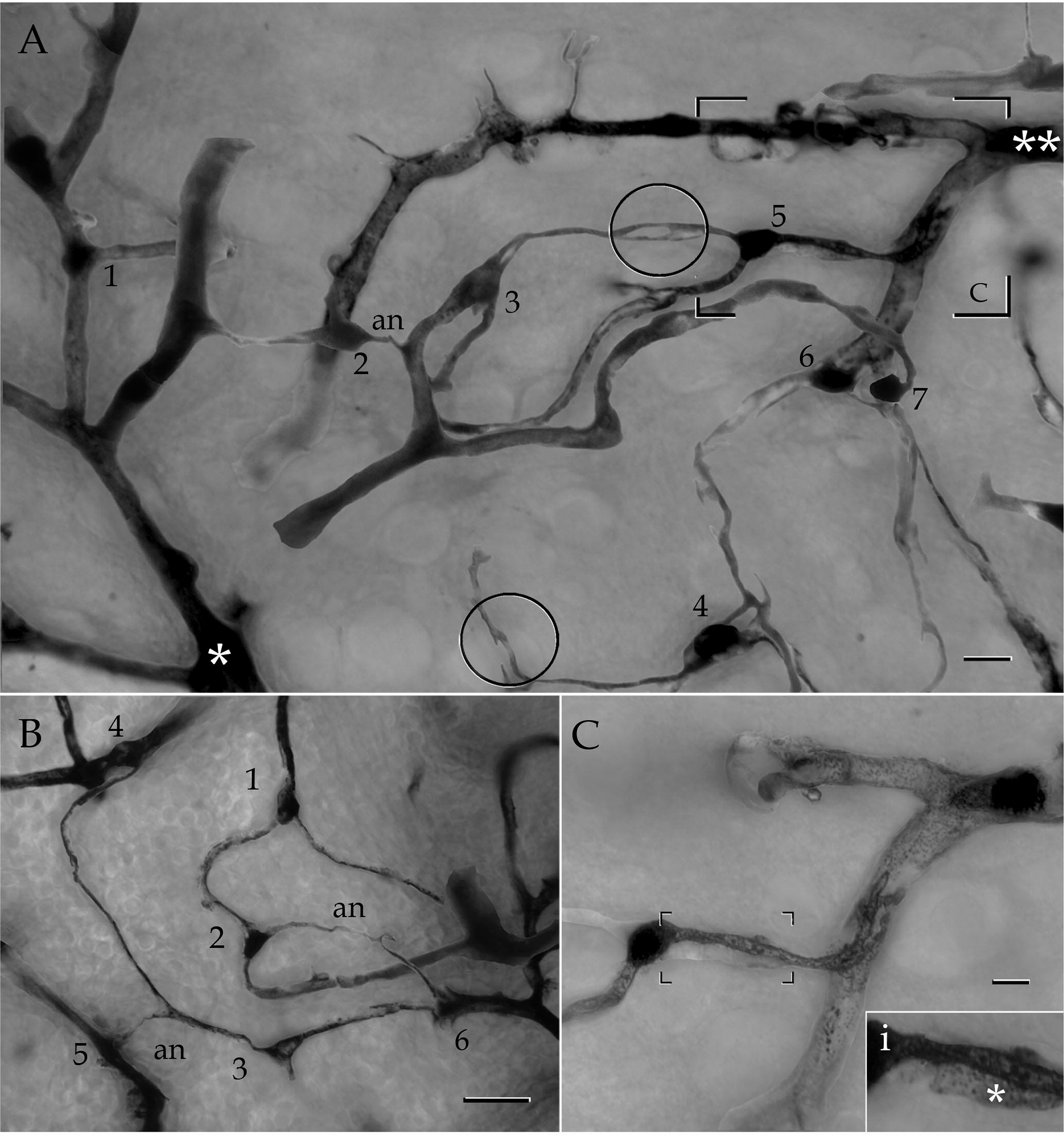
Photomontages of blood vessels and associated pericytes (PCs) in the mouse cerebral cortex. A. Survey micrograph of a capillary bed arising from a penetrating arteriole (asterisk) to converge into a penetrating venule (double asterisk). Seven PCs extend their processes in close apposition to the capillary wall. Notice that processes of most PCs (3-5) form lineal series (see text) whereas others (2) form a tangential shunt as they course devoid a CBV. Two mesh processes (circles) are clearly seen. Hippocampal cortex. B. PCs whose processes organize lineal (1 and 2) and diagonal (2 and 3) interactions. Note that the latter relay upon perforating processes (an) devoid of an associated capillary blood vessel. Hippocampal cortex. an = axonoid. C. Bipolar PC, cell 5 in “A” sending paired primary processes coursing along the capillary wall. i. high magnification view of that part of the PC outlined in “C”. Notice the flat, membranous appendage (asterisk) forming a secondary process. Calibration bars = 30 µm in A, 10 in “B” and “C” ; 5 in “I”.

## Results

As described elsewhere (see Larriva-Sahd et al., 2019) intrinsic arteries of the brain, having medium to large size caliber are encased by a distinct set of cells or perivascular neurons likely to furnish a sensory network. Presently, no PVNs were found in the capillary bed. However, distal to pericapillary arterioles smooth muscle fibers are replaced by pericytes (PCs), so that they form a continuous cellular network surrounding CBVs (Fig. 1). Although thorough descriptions on the structure of the PC are available, a brief description is desirable to provide the context for undescribed characteristics. The nomenclature suggested earlier (Zimmermann, 1923, Sims, 1986) for the PC will be used throughout the text. The soma and processes of the PC visualized in Golgi-stained specimens of the adult rat are summarized in Figures 2A and 3.

**Figure 2.**
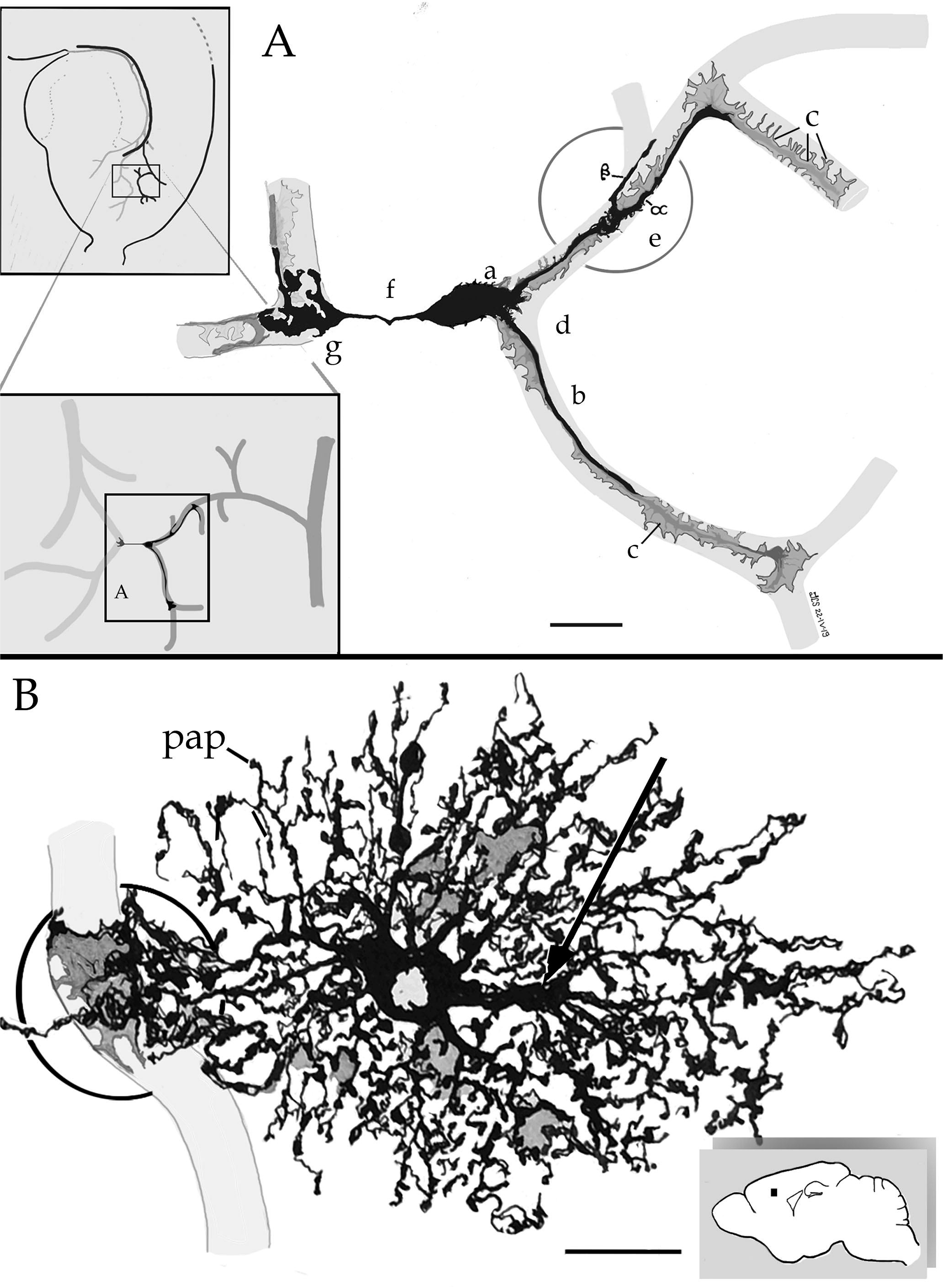
Camera lucida drawings from pericapillary cells and their processes. A. The perikaryon of a pericyte (a) gives rise to three divergent processes, two of them encircling the capillary contour (light gray), and a third one, or axonoid (f), coursing throughout the neuropil to reach an homologue by the terminal vesicular complex (g) . b = primary process; c = secondary process; e = asymmetrical process (AP). Note the two branches structuring the AP, a short branch (β) and a long with membranous outgrowths (α). B. A protoplasmic astrocyte in the cerebral cortex originating, proximal or paraxial-processes (Varela-Echavarría 2017), an end foot (circle) and numerous peripheric astrocytic processes (Reichenbach and Woldburg, 2008). Soft pencil = capillary blood vessel (soft pencil). Scale bars = 3 µm

**Figure 3.**
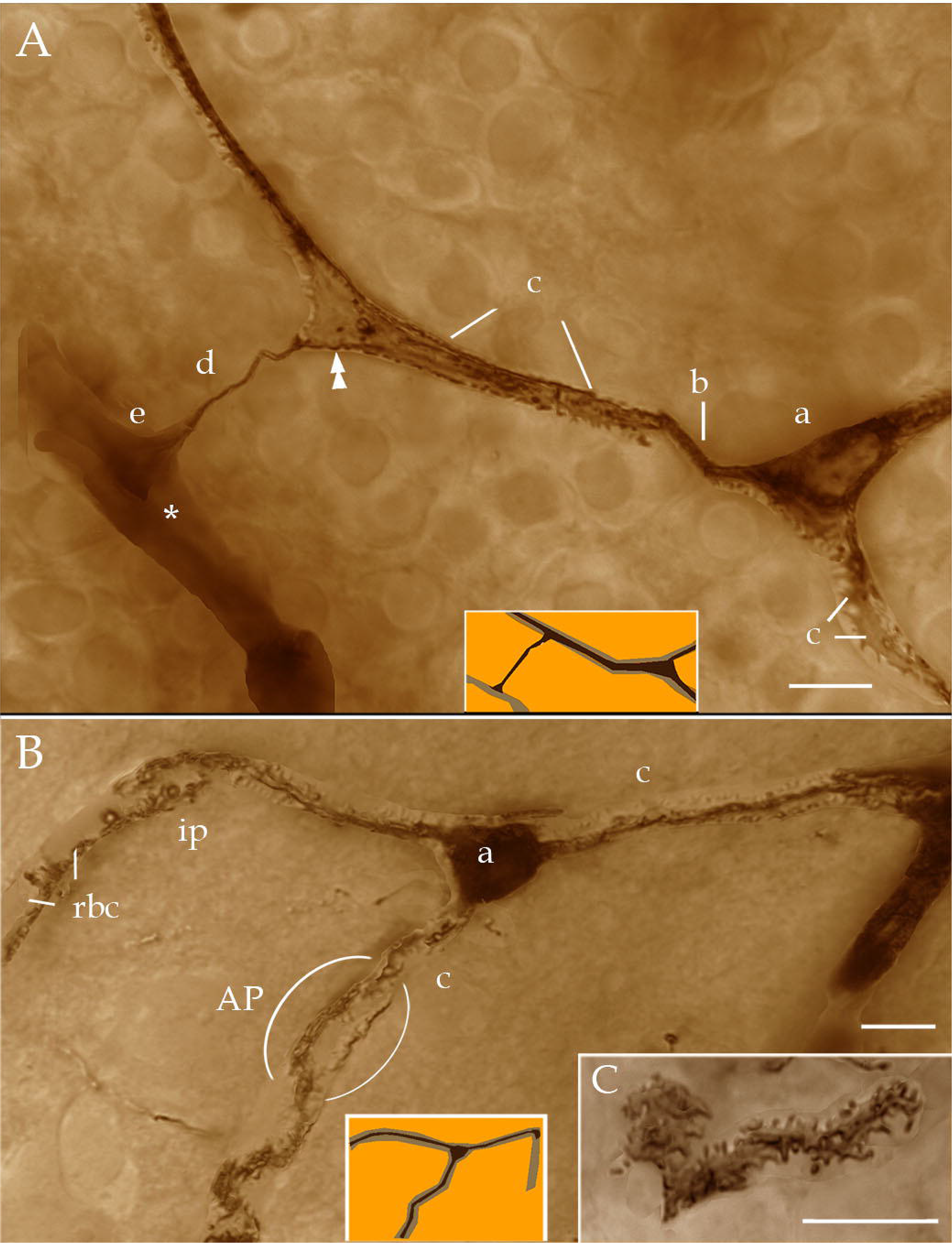
High magnification photomontages of the pericyte and its processes. A. The triangular perikaryon of a pericyte (a) originates divergent processes encircling the unstained capillary wall (light brown-colored in the cartoon). A primary process (b) gives rise to a secondary process (c) which, in turn, sends an orthogonal axonoid (d) piercing the neuropil to form a terminal vesicular complex (e) at the wall of the nearby CBV (asterisk). Notice that the axonoid may be tracked-back to the cytoplasm (double arrowhead) of the parent process. B Photomontages of the pericyte and its processes. Tripolar pericyte (a) riding on a branching capillary blood vessel (CBV). The thick, divergent secondary processes (c) that encircle the external aspect of the unstained CBV which appears lighter containing the ghost of red blood cells (rbc). Ramification of a secondary process originating an asymmetrical process (AP) can be seen (ellipse). C. High magnification, in face view of a secondary processes. Note the short, transverse outgrowths (arrowheads). Scale bars = 5 µm.

PCs form a single row encasing virtually all capillary blood vessels throughout the brain (Sims, 1986; Ushiwata and Ushiki, 1990). Aside of occasional, narrow gaps between them (Berthiaume et al 2018), distal processes from adjacent PCs usually overlap to each other or end foot (EF) (vide infra) (Fig. 4B) When CBV plexuses are tracked from precapillary arterioles to postcapillary venules the extent of the PC covering becomes evident (Fig 1A and B). Alternatively, isolated vessels or small groups of them with diameters of less than 6µm can reliably be considered CBV’s (Figs. 1,3, and 4).

**Figure 4.**
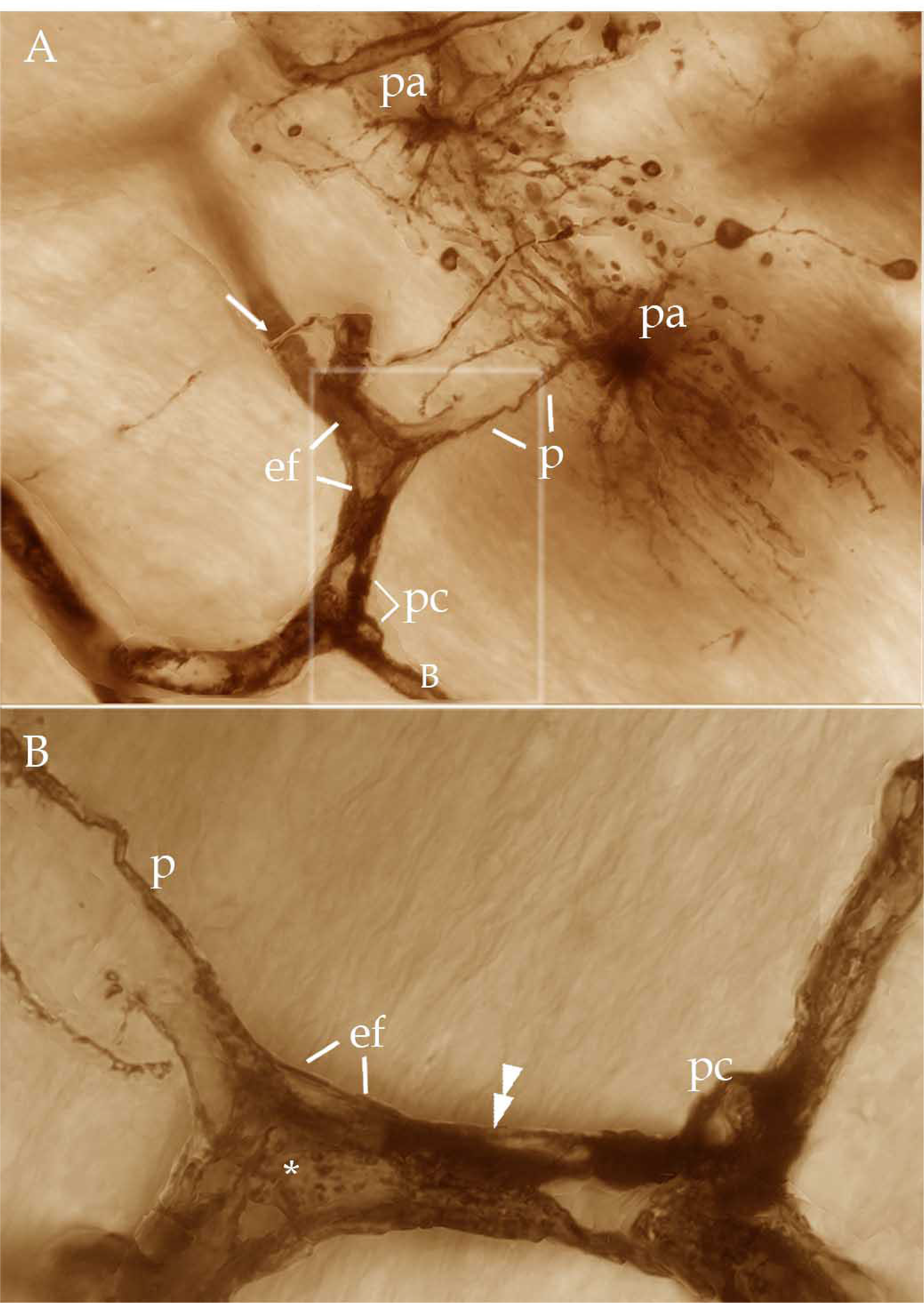
Photomontages from the astrocyte and its processes. A. Protoplasmic astrocytes whose perikaryon (pa) gives rise to numerous radial processes; one of them (p), longer and thicker, evolves to a triangular enlargement or end foot (ef). The latter embraces the capillary wall. B. High magnification view of the end foot framed in “A”. The mantle-like end foot (ef) extends to cover both the ensuing mesh process (asterisk) from a proximal process (double arrowhead) of the nearby pericyte (pc). Scale bars = 5 µm

The pericyte. Characteristics of the PC and its processes as seen in Golgi-stained specimens are shown in Figures 1 to 3, and 4B. The protruding perikaryon of the PC is located either at sites of CBV ramification or in the ensuing capillary shaft, in order of frequency; two or three primary processes arise from the PC cell body (Figs 1 and 2A). These processes are dark and elliptical coursing parallel to the underlaying CBV for lengths that vary from 20 to 200 micrometers. Two types of processes are usually identified in the PC. Namely a dark, argentaffin extension or primary process and a process with light, flat digitations, or secondary process. The former is oval or cylindrical whereas the latter is flat and its finger-like outgrowths exhibit marked pleiomorphism (Figs. 3B and C) recapitulating those observed with scanning electron microscopy in muscular (Mazanet and Franzini-Armstrong, 1982) glandular (Fujiwara and Uehara 1984), or cerebral (Ushiwata and Ushiki, 1990) CBVs. A third distinct development, or “asymmetrical process” (AP) results from the uneven ramification of a primary process (Figs. 1A, 2A, 3B) and, will be further described in later paragraphs. At the electron microscope (Fig. 5) the PC cell body is rounded or triangular and although its contour is generally smooth, numerous short, pleomorphic outgrowths that occasionally overlap with those arising from presumptive neighboring PCs (Fig 5E-G). CBVs gaps devoid of a PC covering are indistinct (Hawkins 2008, Hartman et al., 2014). The PC nucleus bares an overall oval or horseshoe shape whose longest axis adapts to the capillary contour (Fig 5A and C). Like peripheral CBVs, the cerebral PC and its processes remain confined to that area bounded by the inner (iBL) and outer (oBL) leaflets of the capillary basement membrane (Fig. 5-1 A and C) (Burns and Palade 1968). The the so-called fluorescent granular perithelial cell (Mato, et al 1984) described latter on, shares with the PC the area bounded by the inner and outer basal laminae. Characteristics of the PC cell nucleus, and surrounding cytoplasm are presented in panel Figure 5. Briefly, organelle distribution within the perikaryal cytoplasm is asymmetrical (Fig. 5-1A-C). Thus, one pole of the perikaryon is organelle-rich (Fig. 5B-D) containing numerous mitochondria, rough endoplasmic reticulum, and one or two small Golgi apparatus, all of them surrounded by scattered ribosomes and glycogen particles. The opposite PC pole is an organelle-poor area (Fig 5E) whose contour frequently interdigitates with the plasma membrane of a penetrating PC process. Electron microscope series also reveal that the perikaryal profile exhibits numerous in- and out-growths reciprocating processes of a neighboring PC. Similar interweaving interactions are also found in juxtaposed distal processes (vide infra) These membrane-to-membrane interactions are commonly sealed by alternating tight junctions (Burns and Palade, 1968) (Fig 5E-G).

**Figure 5.**
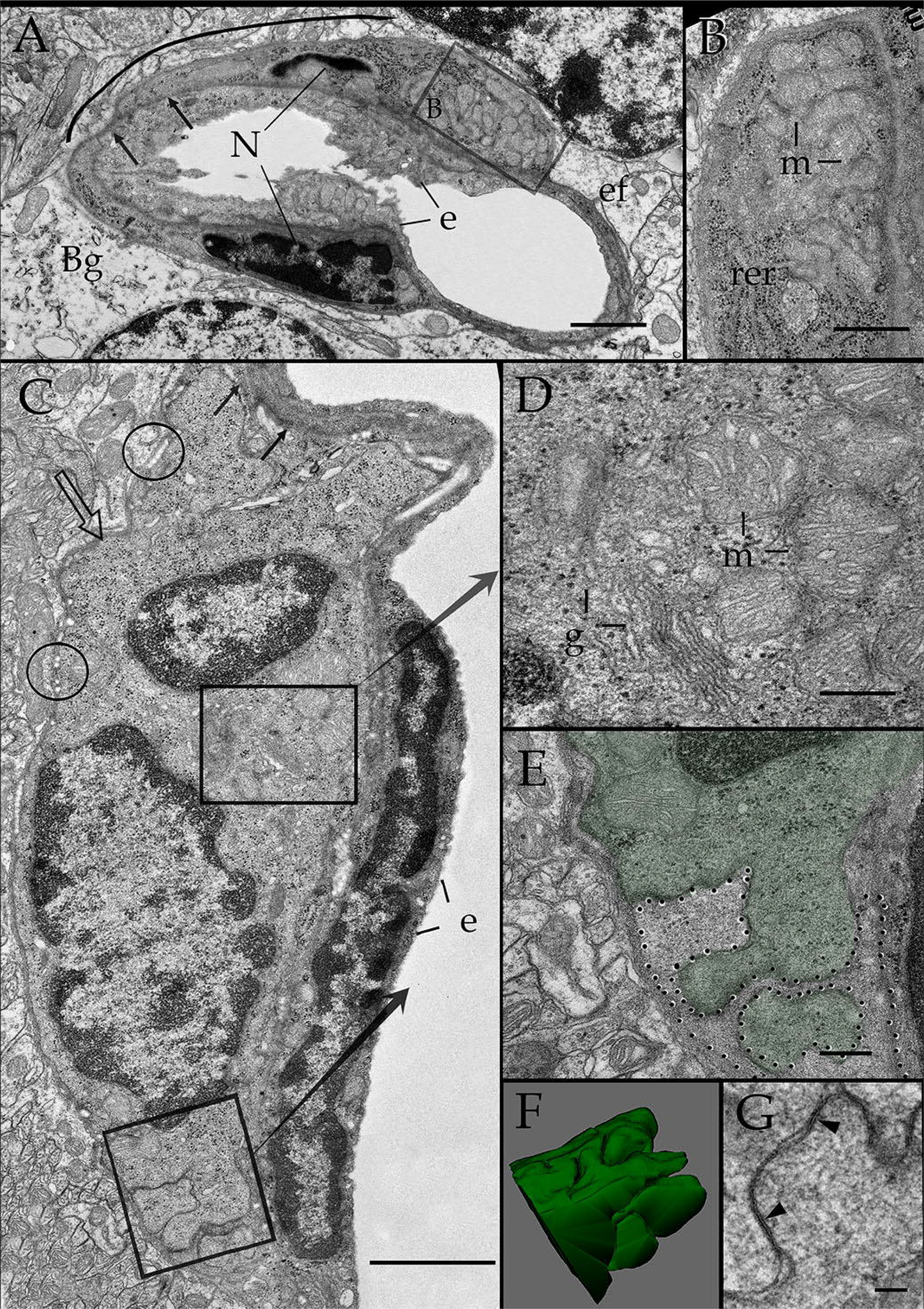
Electron microscopic views of the pericyte (PC): position and interaction with the capillary wall. A. Cross section of the root of two capillary blood vessels (CBVs) partially surrounded by the same PC whose horseshoe-shaped nucleus appears sectioned at either side of the endothelial cell cytoplasm (e). Notice the basal lamina which splits into an inner (black arrows) and outer (hollow arrow) encasing the PC itself. The latter, in turn, faces a Bergman glial cell (Bg) and astrocytic end-feet (ef) forming a long, interrupted covering exposing the outer basal lamina, thereby, open to the neighboring neuropil (curved line). B. Notice the numerous mitochondria and (m) rough endoplasmic reticulum (rer) in a higher magnification view of that area of the perikaryon outlined in “A”. C. Longitudinal section through the capillary wall. Organelles contained by the central (center) and polar (lower) perikaryon of the PC. Notice the inner (black arrows) and outer (hollow arrow) basal lamina containing rows of elastic fibrils (circles) interacting with the neuropil too. D. Cytoplasm of that part of the PC outlined in “C” that contains numerous mitochondria (m) and a distinct Golgi apparatus (g). E. The PC pole (green) delineated in the low part of “C” is composed of in- and out-folds that interdigitate with the distal process of another presumptive PC (coursed by dots). F. 3D view of twenty five sections through the outer aspect of PC karyoplasm. Scale bars = 1 µm in A and C, 0.5 µm in B, D, E, O.2 µm in G.

The protoplasmic astrocyte and its processes (Fig 2B) interact directly with the pericyte. More specifically, the astrocyte gives rise to a variable number of thick paraxial processes (Varela-Echavarría et al., 2017) ending in a cytoplasmic enlargement or EF that encase the capillary wall.

Interaction between pericytes: lineal and tangential linkage. Lineal interactions result from the longitudinal, successive arrangement of PCs along capillary shafts by virtue of junction complexes (vide infra) whereas tangential bridges from a PC process originated from the parent CBV pierce the neuropil to become associated with that PC riding in the nearby CBV

A fourth additional, so far unknown, cytoplasmic specialization of the PC consists of a process that for its resemblance to an axon is heretofore referred to as “axonoid” (AN) (Figs. 2A, 3A and 6). The AN appears to correspond to the solitary PC process identifiable in the illustrations by Zimmermann (1923). Like an axon, the AN is a single, straight process leaving the perikaryon from the apex of a funnel-like extension (Figs. 2A, 6A, C, E, F), or less frequently from a PC process (Figs. 3A, 6B, D). In the latter instance the AN shaft may be identified along the cytoplasm of the parent process (Fig. 3A). Although an AN varies in length (i.e., 15 to 120 µm), it has a uniform diameter (i.e., 0.3 to 0.5 microns), and a zigzag flexure is found into more than two thirds (i.e., 78%) of them (Figs 6A, B, F). Distally, the AN enlarges abruptly to end-up forming a vesicular complex composed of either dense, confluent spherules (Figs 6D, E) or delicate membranous outgrowths (Fig 6B,C,F). The overall area occupied by these complexes is variable and may be as large as 5 microns in the longest axis. Since we cannot find any prior descriptions of this sort of membranous structures, the term vesicular complex is suggested.

**Figure 6.**
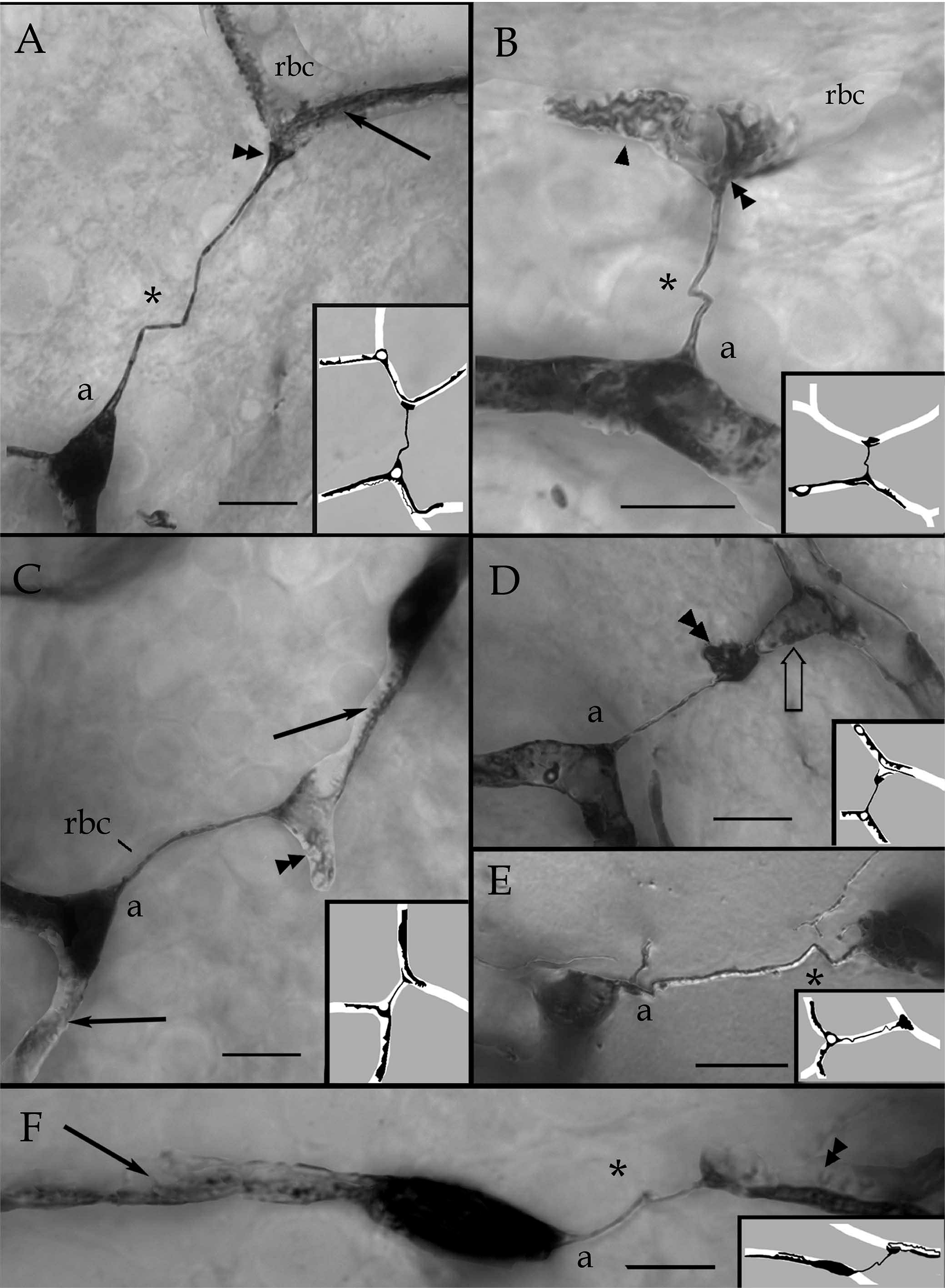
Photomontages of axonoid bearing pericytes (PCs). A. Two capillary blood vessels outlined by the PC perikaryon (lower left) and primary processes (upper right, arrow). Notice that the former extends a perforating axonoid devoid of an associated capillary blood vessel. Notice that the axonoid zigzags (asterisk) at the middle to end into a triangular vesicular complex (double arrowhead), which appose with the primary process (arrow). rbc = red blood cell. B. Another perforating axonoid bridging two capillary blood vessels. The axonoid (a) resolves in a membranous extension (double arrowhead) next to a secondary process (arow head) associated to the capillary blood vessel in the upper part (not shown). rbc = ghost of a red blood cell. C. Two PCs with overlapping, distal processes. Arrows = secondary process, double arrowheads = membranous extension of the axonoid (a) riding in an unstained CBV. rbc = ghost of a red blood cell. D. A perforating axonoid (a) creating a dark-bizarre ending (double arrowhead) next to a mesh process (hollow arrow). E. A PC perikaryon extends an axonid (a) originating a terminal bouton (double arrowhead). Phase-contrast optics. F. A bipolar pericapillary PC whose perforating axonoid (a) resolves next to a secondary process (double arrowhead) in a nearby blood vessel. Arrow = secondary process. The axon appears to terminate at the tip of the dendrite. Double arrowhead = membranous axonal appendage. Scale bars = 3 µm

Tangential interactions: perforating axonoids. Although the association of interlacing PCs (Fig. 3B) paralleling the CBV wall is implicit, a set of ANs eaves the capillary domain to reach processes of a homologous cell riding in a nearby (i.e., different) CBV (Figs. 2A, 6A,B,D,F). These perforating processes were noted earlier in the cerebellum (Zimmermann, 1923) and retina (Williamson, et al. 1980), and are described and found in available series of the human cerebral cortex (Shapson-Coe, et al 2021). Presently, the ubiquity of perforating AN through the forebrain CBVs is evident. Hence, PCs exhibit two different arrangements, namely a lineal or straight, whereby a PC interlaces with its homologous in the same CBV, and transverse or tangential whereby a PC originating perforating AN intertwine with the PC of a different CBVs. Further inspection of the CBV forth and backwards to the tributary penetrating arteriole and vein, respectively, reveals that perforating ANs link PC-CBVs of a heretofore unsuspected asymmetric caliber (Fig 7C, D). It is further noted that parent CBVs linked by perforating ANs arise from distant (i.e., different) tributary blood vessels (Fig 2A). Another thus far neglected feature of perforating ANs is the uninterrupted EF envelope they exhibit (Fig. 7C-D). Observations at the electron microscope confirm first that inter-capillary AN bridges arise from PCs riding in CBVs of different diameter (Fig 8A); second, the PC process form a core in the bridge that is covered by an EF cuff (Figs. 7C,D, 8A-C); third, within the core convergent processes may entangle with each another; and lastly plasma membranes along this cell-to-cell interaction appear to be united by series of tight junctions (Fig. 8C-E), in a similar fashion to that described for the PC perisomatic aspect (Fig 5G).

**Figure 7.**
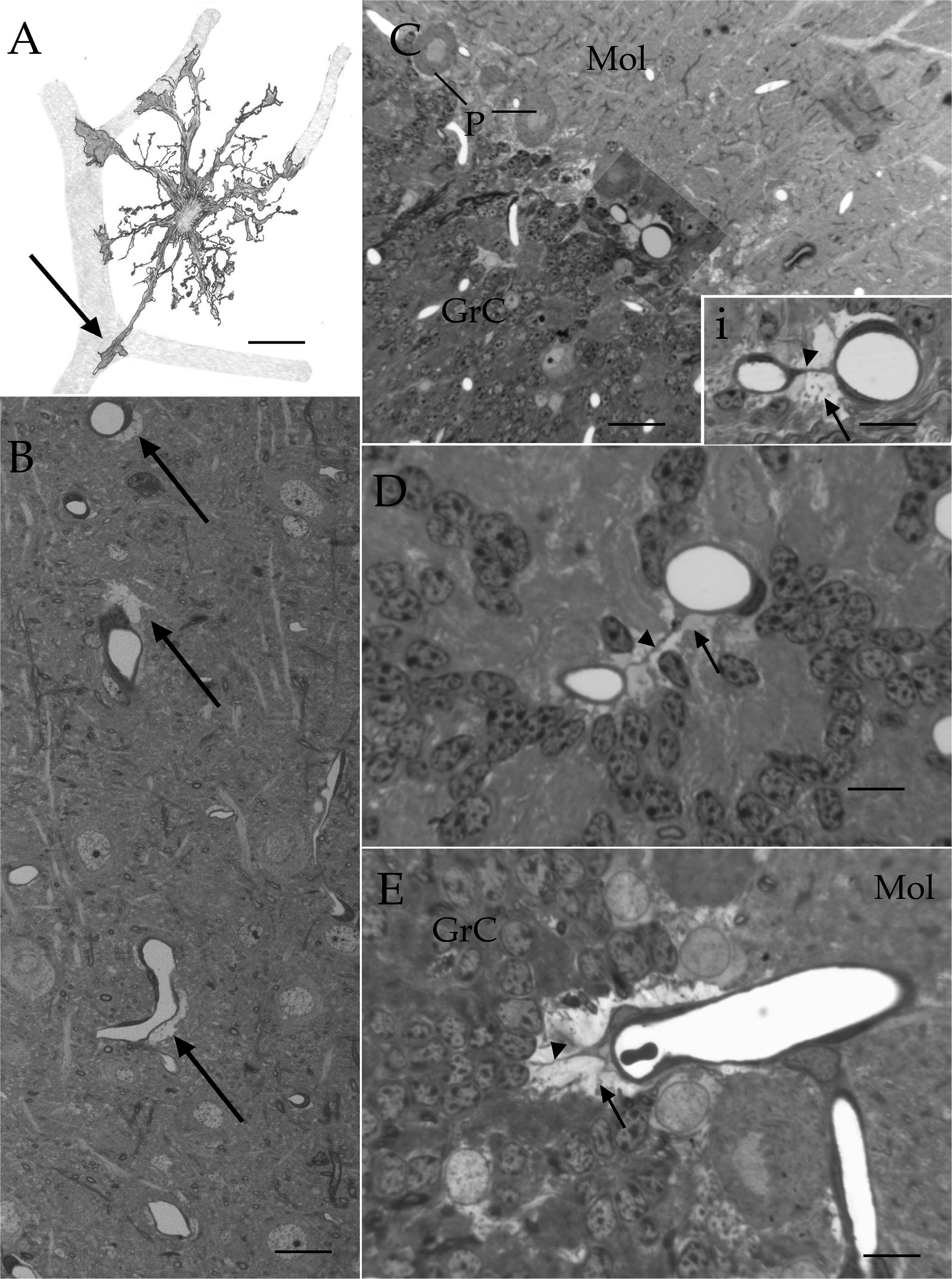
Light microscopic images illustrating the astrocyte and its processes. A. Camera lucida drawing of a protoplasmic astrocyte creating radial processes that terminate into flat end-feet (EF) (arrow) surrounding the adjacent capillary blood vessels (soft pencil). B. Semithin section showing the patch-like appearance of EF (Arrows), firmly apposed to capillary blood vessels. Parietal cortex, toluidine-blue stain. C. Micrograph at the intersection of the molecular (Mol) and inner granular (GrC) layers of the cerebellar cortex; between them there is the row of large Purkinje cells (p). Notice the two capillary blood vessels (shaded) of a first glance dissimilar diameter. i. high magnification of the CBVs in “C” depicting a perforating axonoid (arrowhead) embedded by EF (arrow). D. High magnification micrograph of the cerebellar granule cell layer. E. A tangential section through a capillary blood vessel surrounded by a crescent-like made-up of EF (arrow). A putative PC bride can be seen (arrow-head). Scale bars = 10 µm in A to D; 5 in “i”. Rapid Golgi in “A”; Toluidine blue in C to E. Adult rabbit brain. Scale bars = 5 µm in A, C – E, 10 µm in B, 3 µm in i.

**Figure 8.**
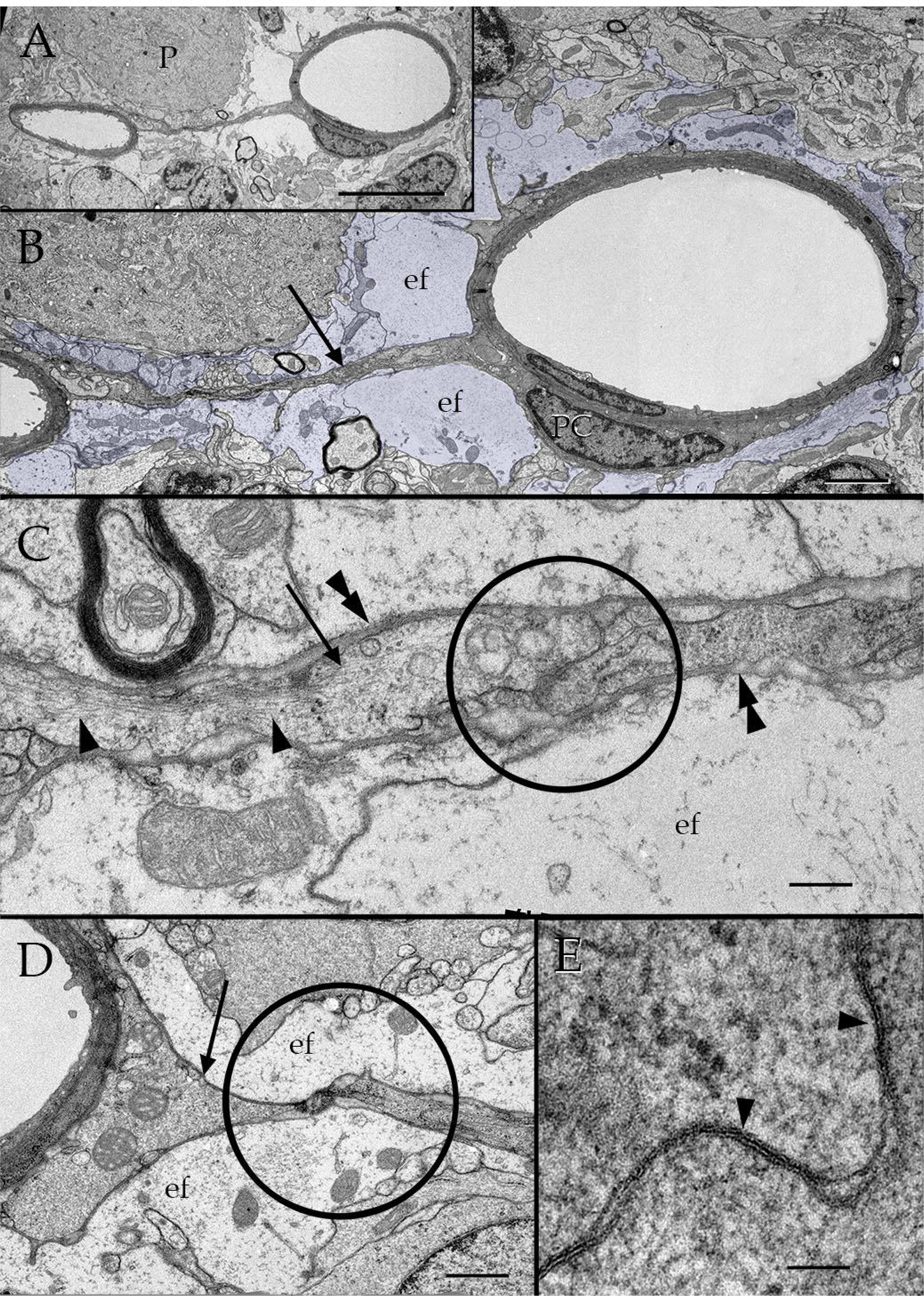
Ultrastructural appearance of the pericytés (PC) intercapillary bridges (ICB). A. Low magnification view of an ICB between two cerebral blood vessels (CBVs). Notice that there is a first-sight asymmetry in vessel caliber. P = Purkinje cell. B. High magnification micrograph from the ICB shown in “A”. Note the complete cytoplasmic cuff (light-blue) that end-feet (ef) paved around both CBV and pericyte processes (arrow). pc = cell nucleus of a pericyte. C. Interaction (circle) between two PC processes (arrows) creating the core of an ICB in the frontal cerebral cortex. Double arrowheads = basal lamina; single arrowheads = microfilament bundles. D. Contact processes (circle) from two PCs structuring the core of an ICB. E. High magnification view of two uniting PC processes. Notice that the processes plasma membranes alternate with zonula occludens (arrow heads). Scale bars = 3 µm in A, 1 µm in B, 0.5 µm in C and D, 0.2 µm in E.

Still another outstanding observation is that the PC perikaryon and its proximal process may be devoid of a CBV (Fig.9). This uncommon extra-capillary location prompts the idea that it corresponds to a neuron rather than a PC. However, one the distal process usually anchors and surrounds a CBV exhibiting the typical location and structure of a PC process. Further, the process leaving the opposite pole of the perikaryon is indistinguishable from a typical AN (Fig. 9). So far only five extra-capillary PC of this sort have been observed (three in the cerebellar granule cell layer and two in the cerebral isocortex.

**Figure 9.**
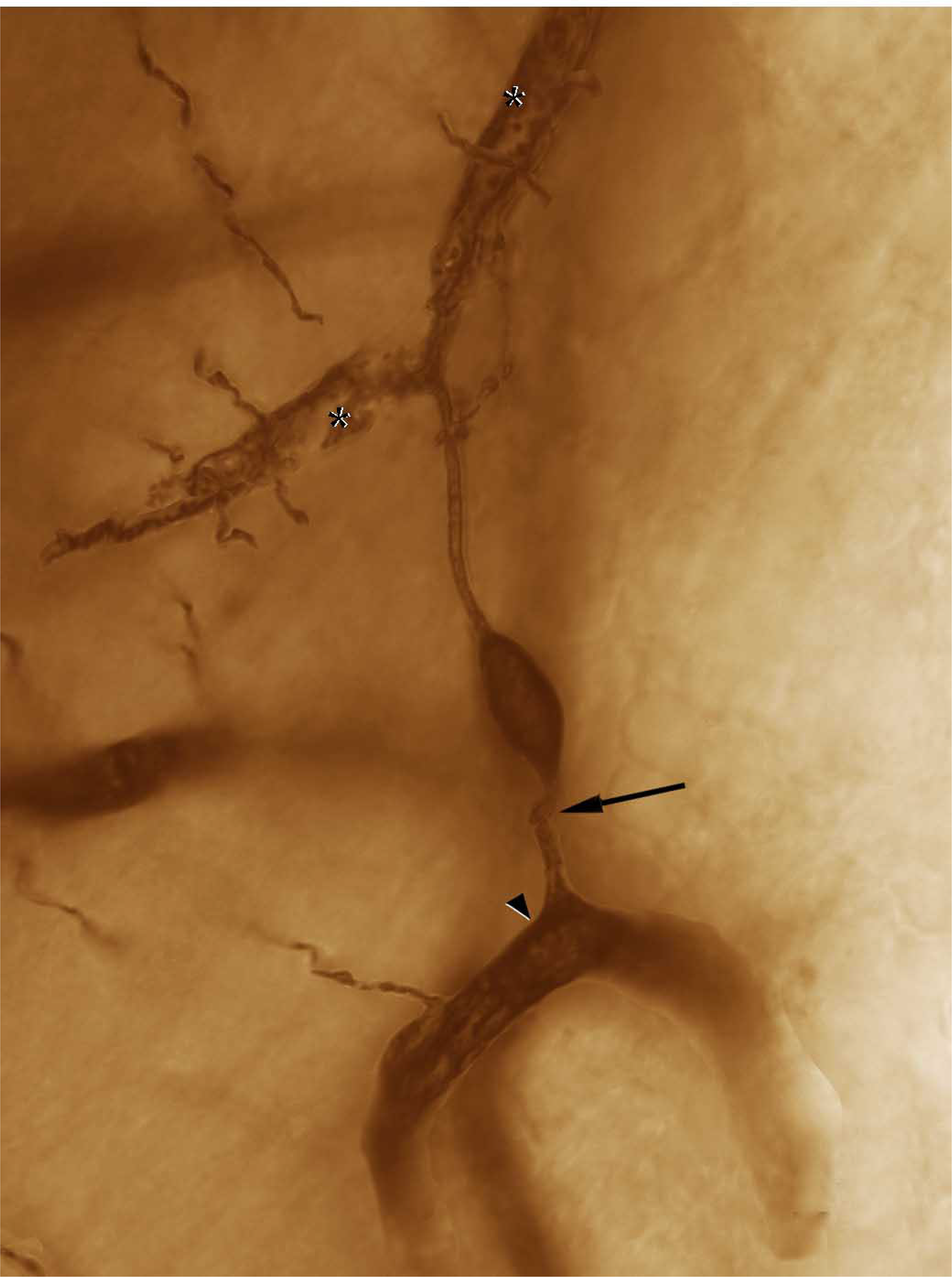
Photomontage of an extra-capillary pericyte. This bipolar PC, surrounded by the neuropil sends an ascending, ramifying processes that gives rise to typical secondary processes (asterisks) entangling a capillary blood vessel (CBV). The opposite pole of the perikaryon extends a distinct axonoid (AN) anchoring in another nearby CBV (arrowhead). Note the discreet flexure (arrow) of the AN beneath the perikaryon. Scale bar = 5µm

Asymmetrical-, neuron-processes, end feet, and perivascular ground substance structure a seemingly distinct organ. A frequent membranous appendage of the PC is that resulting from paired uneven branches or asymmetrical process (AP) (Figs. 2A, 10). Directed search for the AP in various brain areas reveals their ubiquity and adds structural features which we summarized next (Fig 10). First, an AP results from the dichotomous division of a primary process into two different branches termed here alpha (αB) and beta (βB). The resulting horse-shoe arrangement outlines the CBV contour (Figs 10A-C, arrow), being wider at sites of ramification (Fig. 2A,10E double asterisk). The αB shaft stains firmly with silver and it measures one to two microns in diameter, although its length is highly variable measuring 20 microns or more. The αB stands out for its four to seven thin, mantle-like, outgrowths which occupy the concavity of the AP. These distinct formations provide to the αB an overall appearance to a shelf with graceful ornaments on it (Figs 3A, 10). The βB is thinner, shorter and darker than the αB, ranging from 0.5 to 1.0 microns in dimeter and 5 to 15 microns in length. An important landmark for αB identification is the pale, lucid appearance as an extension of the basal lamina filling up the AP concavity (Fig. 10) Specimens from various mammals, reveal conservation of PCs in mouse, albino rat, Guinea pig, kitten, rabbit, Rhesus monkey, and human brain conservation (Fig. 11). At the electron microscope (Fig. 12), the αB-AP lies underneath the EF and its βB just outside the outer basal lamina, thereby, paralleling the CBV lumen (vide infra). Serial sections and 3D reconstructs at the root of a CBV provide further characteristic of the AP. To start with, is the fine structure of the basal lamina filling up the αB concavity. This consists of a two-fold basal lamina (Fig. 12D and E); namely, a sharp layer of moderate to high electron-density, or *pars densa*, associated with the βB, and an electron-diaphanous one, or *pars lucida*, surrounding the membranous outgrowths of the αB. An additional distinct feature of the *pars lucida* is that it its a homogeneous matrix is traversed by numerous, strands from 15 to 18 nm thick which extend radially, between αB and βB inner aspects (Fig. 12E), mimicking those embedded in the Pacinian corpuscle basal lamina (Munger et al., 1988, Dubovy and Bednarova 1999). 3D reconstructs of the αB and βB (Fig.12B and C) reveal the unique interaction between them. Tiny, inconspicuous axons piercing the EF observed in single sections (Fig 12A, arrows), correspond to a set of synaptic outgrowths termed here “golf club-like extensions” (GCLs) (vide infra) reveled by 3D reconstructs (Fig. 12B and C).

**Figure 10.**
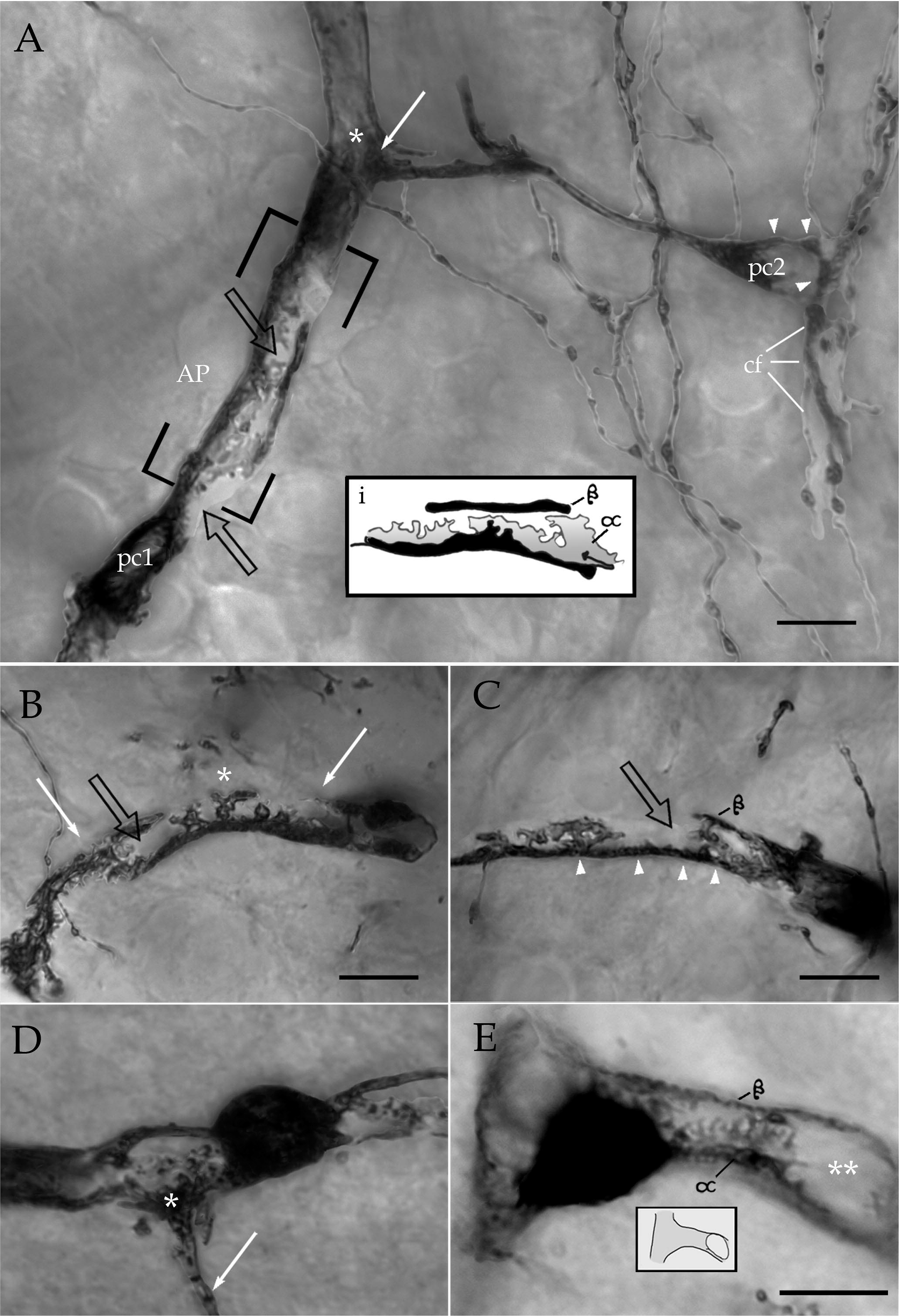
Membranous specializations of the pericyte (PCs). A. Anastomosing (arrow) capillaries surrounded by two PCs in the rat cerebral cortex. One of them (pc1) extends a primary process originating an asymmetric process (AP). It is evident the irregular membranous outgrowths arising from the thicker, or alpha arm (α) towards the shorter, thinner, or beta arm (β). Notice the diaphanous basal lamina (hollow arrows) embedding the outgrowths of the alpha arm. Another perikaryon (pc2) sends a horizontal axonoid whose terminal vesicular complex (arrow) overlaps distal process (asterisk) of the pc1. cf = climbing fiber encircling (arrowheads) the perikaryon and proximal processes of the pc1 i. Camera lucida drawing showing the asymmetry between the alpha (α) and beta (β) processes organizing the AP shown in “A” with outgrowths of the alpha branch occupying the recess between the two branches. B. An AP (between arrows) surrounding an unstained capillary blood vessel and an emerging branch (asterisk). Hamster posterior thalamus. C. A long AP in a capillary blood vessel of the rabbit frontal cerebral cortex. Notice here and in “A” and “B”, the pale, chromophobic area (arrow heads) embedding the membranous outgrowths from the thick argentaffin, alpha processes, which oppose to the beta (β) process. D. Specimen of the mouse cerebral cortex showing the perikaryon of a pericyte bounded by two APs. Notice that the arriving terminal vesicular complex (asterisk) of a penetrating axonoid to the AP domain. Arrow = axonoid. Cerebral cortex. Perikaryon surrounding a capillary blood vessel ramification. Notice that the cell gives rise to a AP along the capillary shaft (right). E. Rhesus monkey frontal cortex of a PC whose proximal processes organize an AP that, distally encircling an arising capillary blood vessel (asterisks). Calibration bars = 5µm.

**Figure 11.**
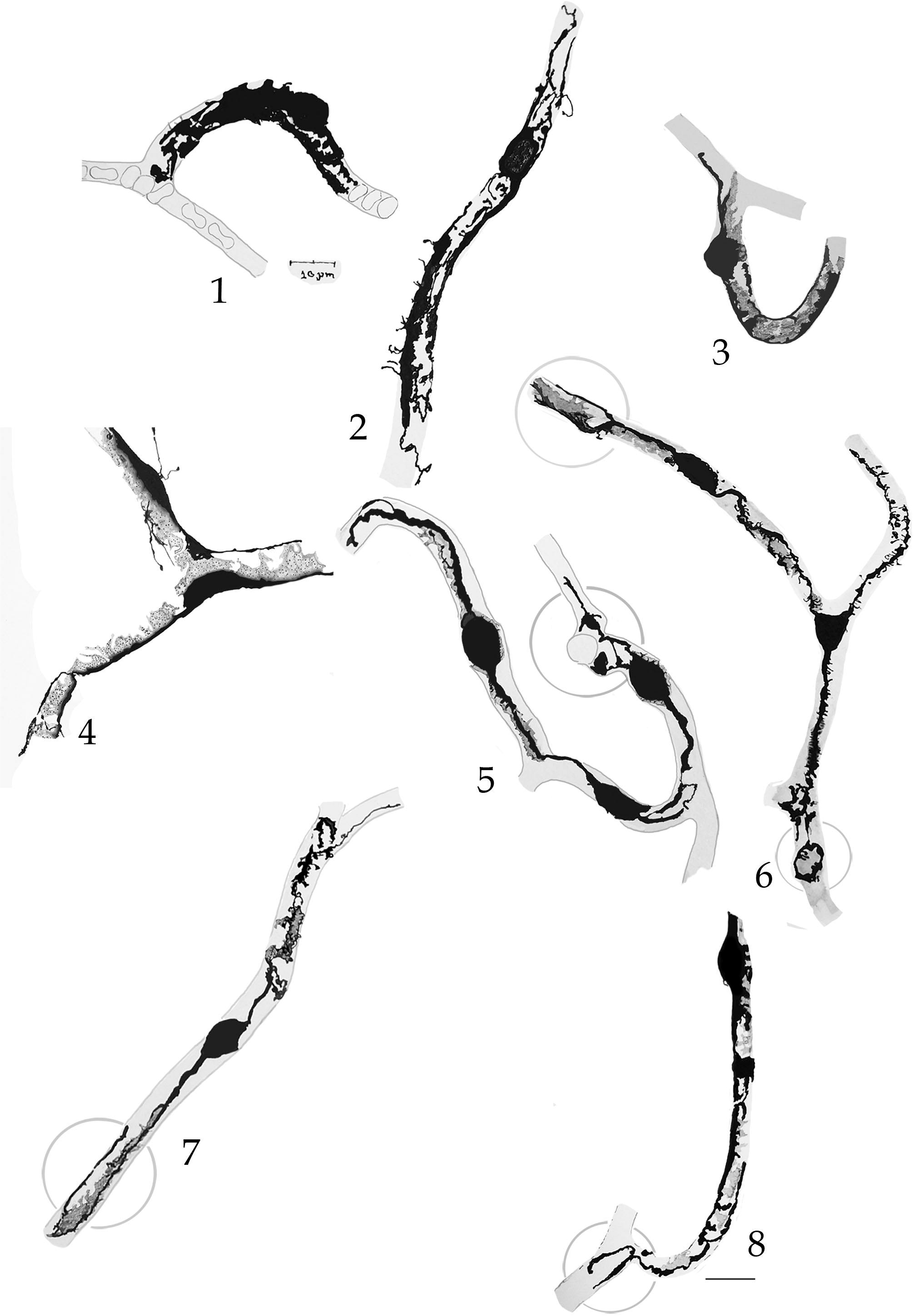
Camera lucida drawings of the pericyte in capillary blood vessels of various mammals. Human (cells 1 and 2), rhesus monkey (cell 3); cat (cell 4), Guinea pig (cell 5), rabbit (cell 6), mouse (7) and rat (8). Circle = mesh process. Cerebral cortex, rapid Golgi technique. Calibration bar 5 µm.

**Figure 12.**
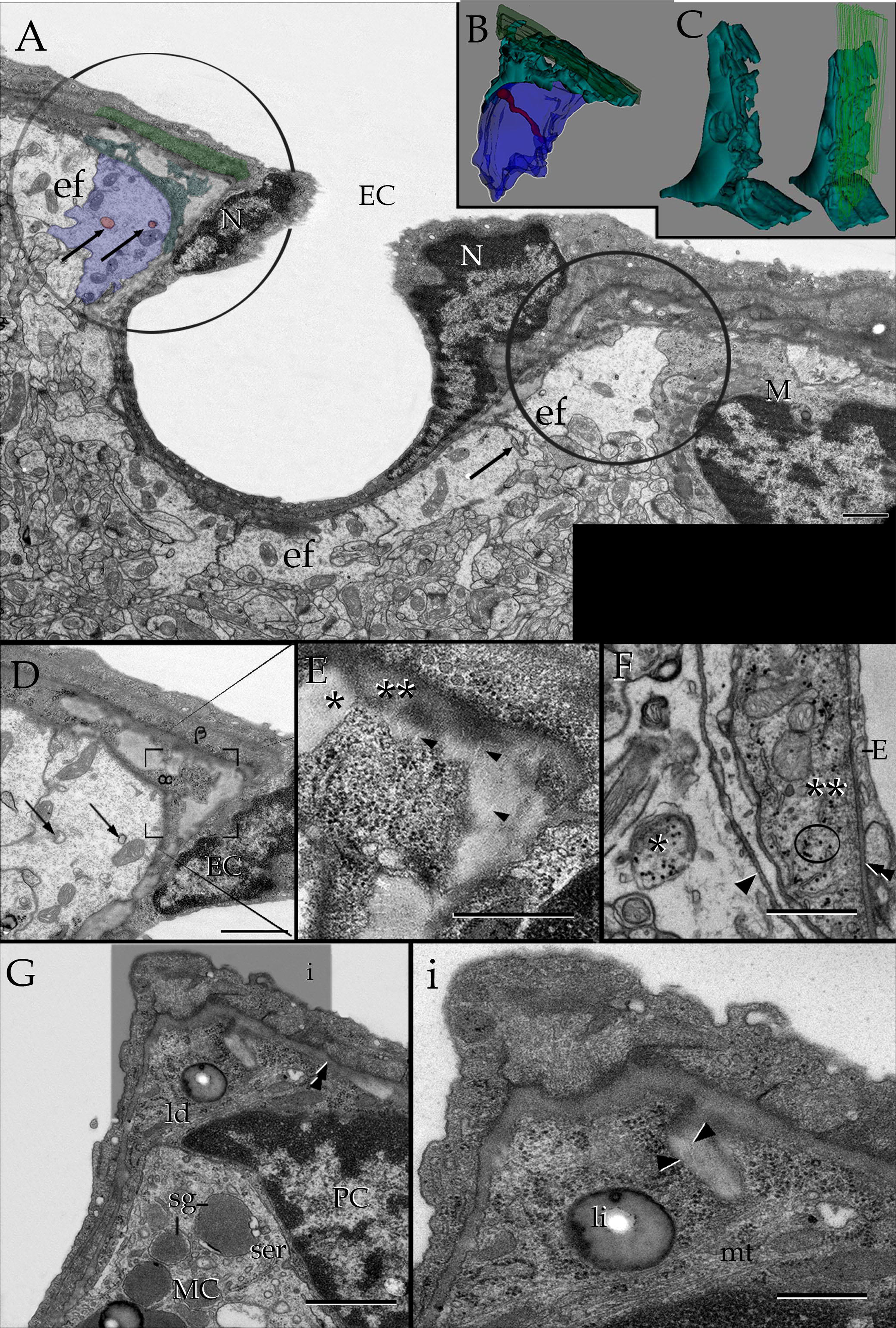
Electron microscopic micrographs of the capillary neurovascular unit in two- and 3D images. A. Survey micrograph at the site of origin of a transversely sectioned capillary blood vessel. The AP (green) consisting of paired (α and β) branches (arrows). Because of the annular shape of both endothelial cell nucleus (N) and the AP proper (circles), they can be seen twice at either side of the capillary root. Astrocytic end-feet (ef), and a nearby microglia (M) provide a continuous covering to the AP. Notice that the end-foot (blue) covering the AP proper, is pierced by a thin nerve (red, arrows), or golf-club-like terminal. B. Reconstruction of fifty-five consecutive sections through that part of the AP encircled in “A”, left side. The external structure of the asymmetrical, opposing α (turquois) and β(green-colored) arms of the AP is evident (see Figs. 2 and 10). Notice that the outer aspect of the AP is covered by the end foot (blue) that, in turn, is traversed by the golf club-like fibril (red) extending a distinct enlargement next to the AP. C. appearance of that part of the alpha component (left) of the mesh processes facing the beta outgrowth that appears outlined in light green (right). D. Micrograph through the opposing alpha (α) and beta (β) arms of the AP process reconstructed in “B and C”. The stem nerve originating the GCL reconstructed in “B” can be seen (arrows). E. High magnification micrograph from that part of the field labeled in “D”. Note the basal lamina embedding the AP exhibiting two, distinct constituents: a thick, electron-opaque, component (double asterisk) or pars densa and an electron-diaphanous or pars lucida (asterisk), the later is pierced by light, radial threads (arrowheads). F. Section through the wall of capillary blood vessel whose endothelium (E) covers a pericyte process (double asterisk) containing abundant electron-opaque glycogen particles (circle), mitochondrial and sparse rough endoplasmic reticulum. Single arrowhead: outer capillary basal lamina; double arrowhead = inner capillary basal lamina. G. A section through an endothelial cushion protruding to the lumen of a capillary blood vessel. Underneath the inner capillary basal lamina (double arrowhead), there are portions of a pericyte (PC) whose cytoplasm contains numerous free ribosomes, scarce rough endoplasmic reticulum, and a solitary granule with lipid droplet material (li). Next to the latter there is the weakly electron-dense cytoplasm of a perithelial fluorescent cell (MC), containing large, rounded, secretory granules (sg) surrounded by cisternae of the smooth endoplasmic-(ser), and rough endoplasmic-reticulum. i. High magnification view depicting an invagination containing the electron-lucid basal lamina associated with a AP as confirmed by identification through the adjacent sections. Note the presence of light, radial fibrils (arrowhead) embedded in the pars lucida bounded by the invagination itself. mt = microtubules. Calibration bars = 1µm

A singular feature of the AP is its frequent association with the endothelial cell, end-foot, and Mato cell; although, in no case are these interactions direct because the inner and outer basal laminae interpose (Figs. 12E, G, i) between them. A distinct set of nerves arising from the perivascular neuropil invaginate through the EF to settle on that part of it overlaying the AP proper (Fig. 12A, B). Because of the long shaft originating an end dilation of these nerves signify the GCLs just mentioned above. The source and fine structure of GCL and its interactions with the CBV will be a core subject of the following paragraphs. However before proceeding with the description of the pericapillary neuropil, it is necessary to highlight the structure of the second cell type, or Mato cell, associated with the capillary wall.

Mato cell. The ubiquitous perithelial fluorescent cell PFC (Mato et al, 1980, 1984), also termed granular pericyte (Mato et al 1996) or, later on type II mast cell (Dimitriadou et al., 1984,Yang et al., 1996), shares with PCs the intermembranal compartment between the iBL and oBL (Fig. 13, 14) (Graeber and Streit, 1990). Because of this, both are considered to be an integral part of the NVU (Ribatti 2015; Li, et al 2017). Thus, the rounded PFC soma lies next to CBVs (Figs. 12G, 14), usually at sites of ramification (Fig. 13A) detected in a decreasing frequency in the thalamus, basal ganglia, olfactory bulb, hippocampus, and isocortex. The perikaryal PFC contains an oval nucleus surrounded by numerous granules larger that 1µm providing an overall foamy appearance to the cell. Like a PC, the Mato cell sends long paired cytoplasmic processes alongside CBV’s wall. Unlike the former, PFC processes may be exceedingly thin (Fig 10A) and equally longer than100 microns. The presence of solitary granules surrounded by the same organelles contained by the perikaryon (Fig. 13) secure identification of distal PFC processes (Fig 13E). Another cytoplasmic landmark of the PFC is the large area occupied by the smooth endoplasmic reticulum surrounded by anastomosing granules (13A, 14) whose limiting membrane is reinforced by a fussy material. A set of the later granules, scattered next to the plasma membrane, contains distinct electron-diaphanous material seemingly identical to the collagen fibrils scattered in the basal lamina. Identification of continuity between cytoplamic granule-contents and extracellular collagen fibrils (Starborg et al., 2008) in series (Fig. 14) holds an emerging notion favoring the PFC role in basal lamina synthesis.

**Figure 13.**
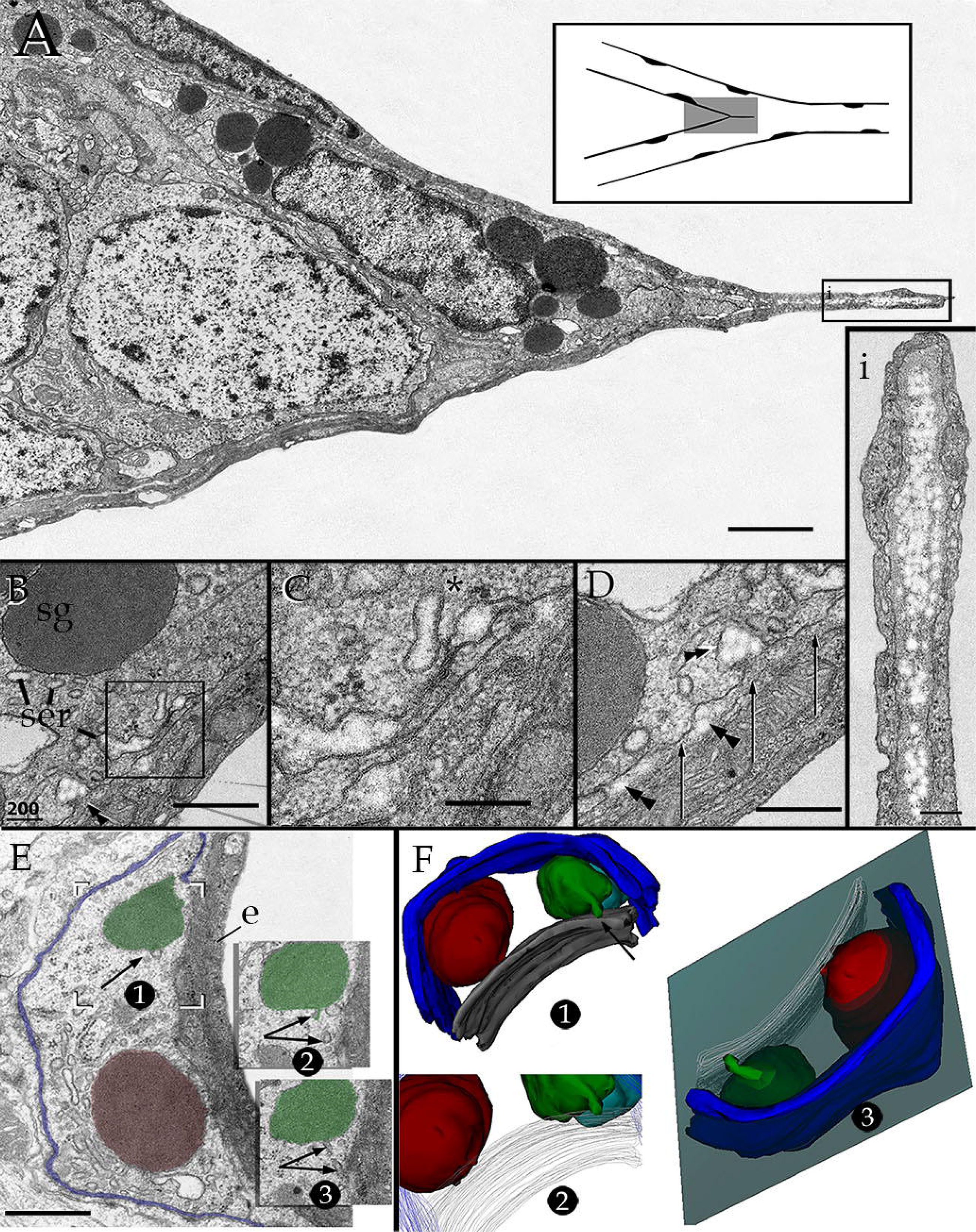
Bi- and 3-D electron microscopy of the perithelial fluorescent cell (PFC). A. A PFC at the mesangium of a forked capillary blood vessel (CBV) which contains numerous electron-dense, secretory granules (sg) scattered in the cytoplasm. An endothelial shaft protruding from the mesangium to the lumen can be seen. i. High magnification view to depict the core of the shaft containing electron-lucid clusters of collagen fibrils. B. Micrograph of the interaction of the PFC cytoplasm with the extracellular matrix, notice the anastomotic tubes of the smooth endoplasmic reticulum (ser), containing electron-lucid material. Extrusion of the electron-lucid material to the endothelial basal lamina (double arrowhead) may also be seen. C. High magnification view of the area outlined in “B” whereby the appearance of ser content is further appreciated. D. Micrograph contiguous to that clusters of collagen fibrils are partially encircled by the smooth endoplasmic reticulum (upper right) or embedded (arrowheads) in the endothelial inner basal lamina (arrows). E. Series throughout distal processes of the PFC. In micrographs 1 to 3, a tiny duct (arrow) between the secretory granule (green) and extracellular matrix is shown. F. 3-D views of the series shown in “E”. 1. Upper view of secretory granules. 1. and 2. The continuity of the narrow tube (arrow) with the secretory granule (green) and endothelial basal lamina (gray) is evident. Blue = external basal lamina. 3. Like the perikaryon (B to D), the process of the PFC is encased by a continuous basal lamina formed by the endothelial (gray) and external (blue) capillary basal laminae. Calibration bars = 1 µm in A; 0.5 in B – E; 0.2 in i.

**Figure 14.**
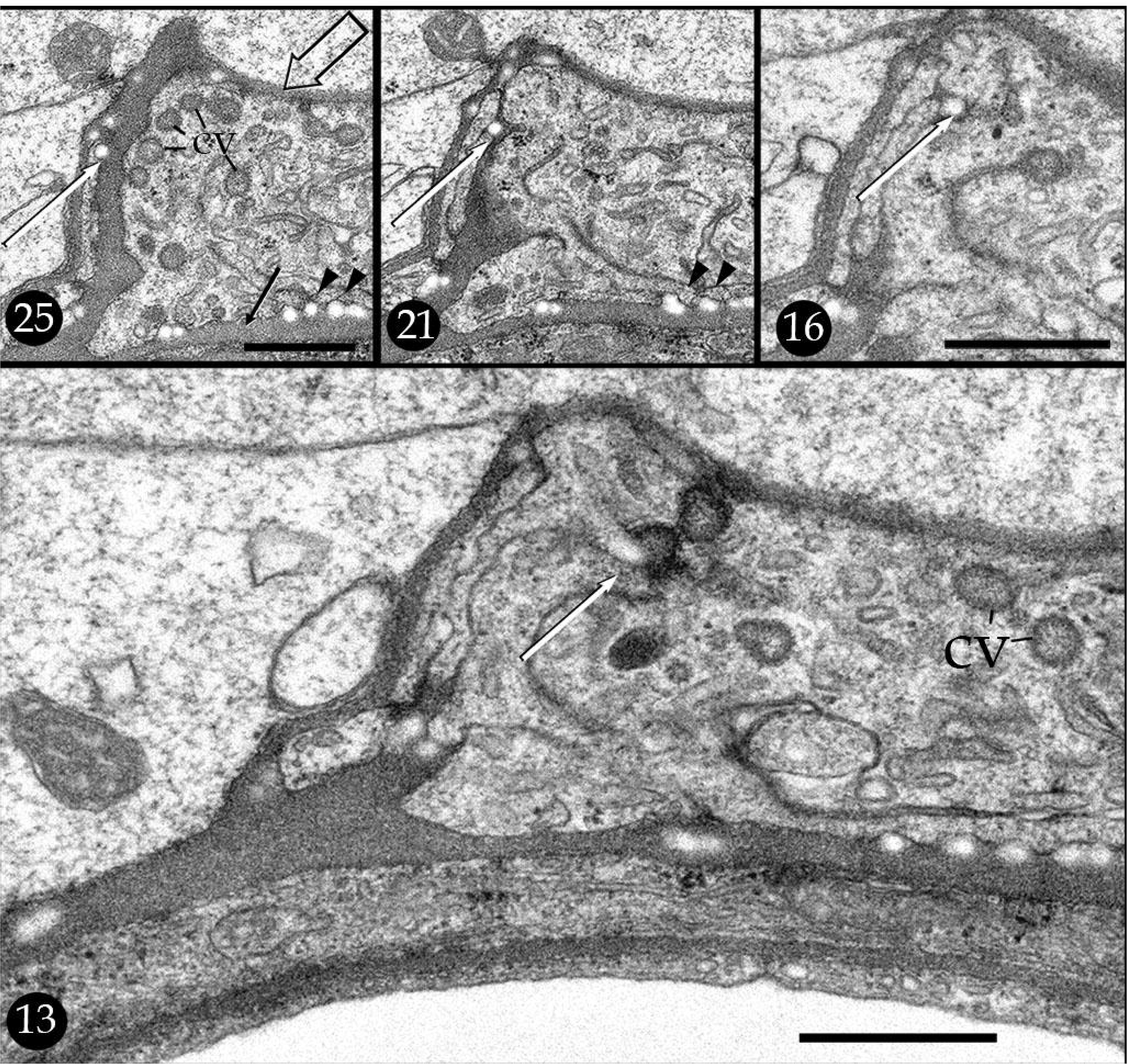
Semi-serial sections (Arabic numbers) of the juxta-nuclear cytoplasm of a Mato cell. Notice in section 25 that the later lies between inner-(black arrow) and outer (hollow arrow) basal lamina which embed clusters of cross-sectioned collagen fibers (arrowheads). The series shows the progression of a collagen fibril (white arrow) from being embedded in the outer basal lamina (white arrow) (sections 25 and 21) to appear invaginated (section 16) and, then, to the lumen of secretory vesicle, (white arrow, section 13). cv = coated vesicles. Scale bars = 0.5 µm

End-feet and pericapillary tripartite synapses. Although few uncommon extensions EF lie between the oBL-iBL area, EF rests on the oBL alternating with occasional Microglia (Graeber and Streit, 1990) or scattered throughout the neuropil (Varela-Echavarría et al 2017). Because the contribution of the microglia to the pericapillary structures described here is absent or occasional (Fig. 12A), microglial cells will not be considered here.

The EF mediates between the pericapillary neuropil and CBV proper. The abluminal surface of the CBV is partially covered (Fig. 5 A, C, 15A) (Bertossi, et al 1997) by a set of EF (Figs. 2B, 7A, 12A), which adapt to the capillary wall and adjoining neuropil in an inseparable fashion (Figs. 15-18). Under the light microscope the EF consists of a pale, chromophobic enlargement of a parent process encasing the capillary wall. Perivascular EF location is by and large the most common, however a set them extend from the CBV wall to cover PC inter-capillary bridges whose core consists of presumptive ANs (Figs. 7C-E, 8A-C). In either case, the interaction EF-AN is indirect because the oBL interposes between them (Figs. 8C,D). As shown in Figure 15, each EF covering the cortical or thalamic CBV wall displays at least three distinctive portions. A first part is a highly convoluted or, convex where the EF interacts directly with surrounding neuropil. This part is continuous with the EF pedicle (Figs. 2B, 7A). A second, or inner part (i.e., capillary) is smooth and concave as it outlines the oBL. Lastly, is a lateral, interdigitating part that reciprocates that of the neighboring EF with whom defines distinct gap junctions (Fig 15B)(Bertossi, et al 1997). Overall, EF stand out from both the neuropil and CBV proper by their low electron-lucid cytoplasm having a paucity of organelles although distinct clusters of mitochondria and smooth- and rough endoplasmic reticulum may none-the-less be seen next to the oBL (Figs. 15C, 17D, E). Other than the later, we could not identify the florid collections of secretory organelles described elsewhere (see Boulay, et al, 2017). A defining feature of the EF is that it is virtually devoid of bundles of intermediate filaments which otherwise may easily be seen in the perikaryal or paraxial, proximal processes cytoplasm (Hama et al., 1994, Varela-Echavarría, et al 2017). An additional, distinct characteristic of EF is that its plasma membrane defines a continuous envelope to the intended nerves interacting with the capillary wall (Figs. 15E, F, I, 16A, E-G) (see below). To emphasize, once a prospective pericapillary nerve approaches the parenchymal convexity of the EF, it proceeds towards the capillary basal lamina engulfed by the cell membrane of the later; hence when the penetrating nerve reaches the oBL, the inner and outer EF cell membranes interpose between them (Fig. 17 A, B). Alternatively, PCNs also approach the oBL *between* two EFs remaining sequestered thereby preventing a direct contact with the oBL. Aside of either modality of EF envelope, nerve processes do not contact the oBL (Fig 17A-D). Still another novel EF characteristic takes place at its vascular aspect. There are one or two long invaginations (Fig. 15D, E) left by a longitudinal outgrowth of the oBL, described in the following accounts. In precis, the EF surface exhibits the impressions left by penetrating elements at the superficial and deep facets of it (Fig. 15D-F).

**Figure 15.**
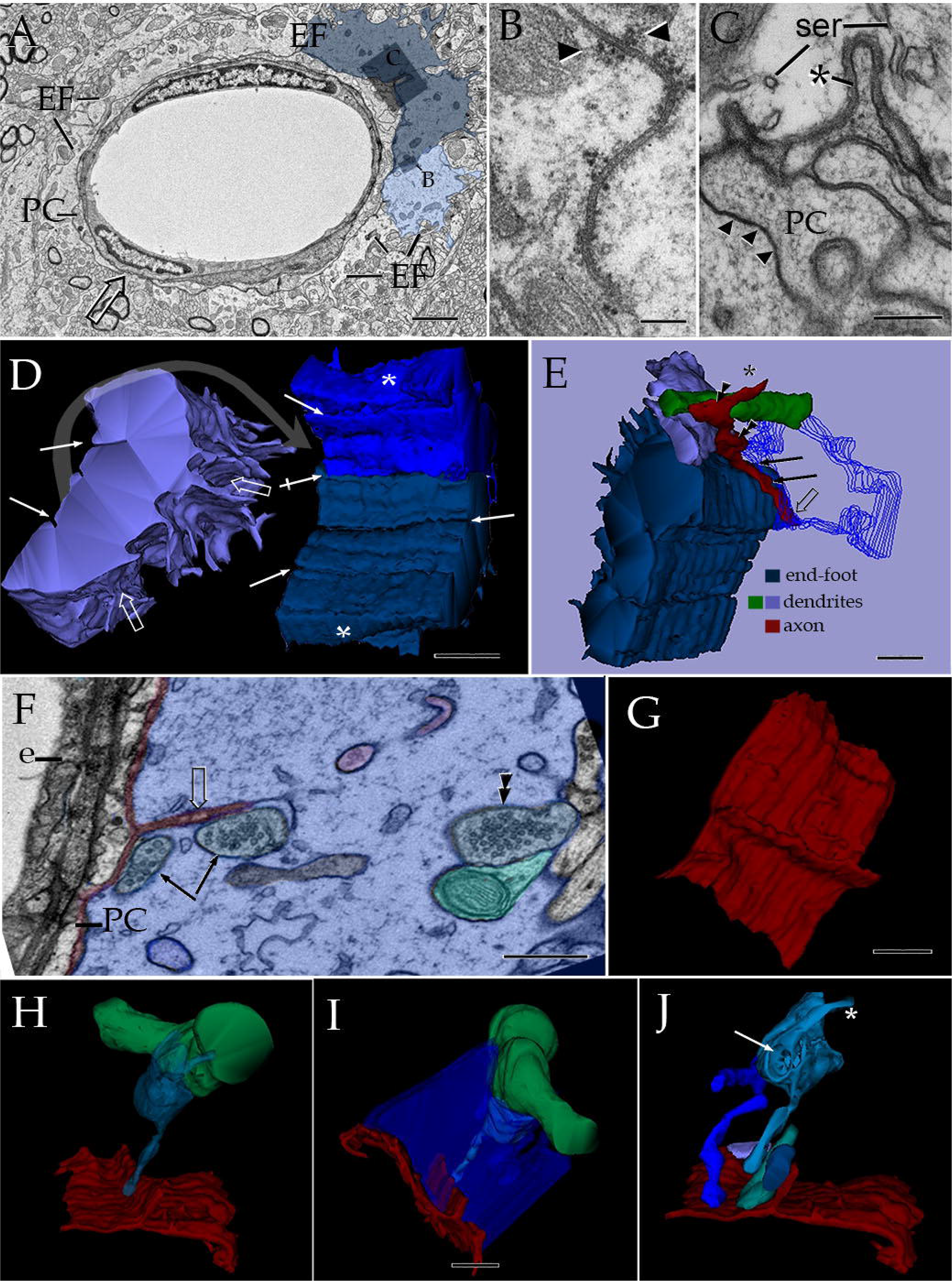
Electron microscopic inner structure and external appearance of glial and neural specializations of the pericapillary domain. A. Survey micrograph of a capillary blood vessel in the cerebral cortex from the same specimen than that shown in Fig. 7B, upper arrow. Notice the discontinuous (hollow arrow) mantle of astrocytic end-feet (EF) built up between the capillary wall and the neighboring neuropil. Two end-feet shaded in pale blue are featured in panel figures “D” and “E”. B. Intersection of two end foot sealed by a long gap junction and clusters of glycogen particles (arroheads). C. A longitudinal crest (LC)(asterisk) of the external basal lamina. Underneath there are interlacing pericyte processes (PC) protruding to the LC and united by tight junctions (arrowheads). C. High magnification view of two presumptive interlacing processes (PC) bounded by the outer basal lamina. Asterisk = longitudinal crests containing protruding processes. Arrow heads = tight junctions between processes. ser = smooth endoplasmic reticulum. D. 3D reconstruction from one-hundred and fifty sections. The elaborated, convoluted parenchymal surface (left, hollow arrows) of the end-foot contrasts with the concave, smooth appearance of the inner, stromal aspect (right) of the adjacent end-feet showing longitudinal sulci (white arrows) that the LC imprint. Asterisk = lateral aspect of the EF. E. Complete (dark blue) and incomplete (outlined in light blue) reconstruction of the end feet shown in “D” to disclose the neural processes between them. An arriving axon (asterisk) gives rise to a modified tripartite synapse (double arrowheads) . The axon fibril (asterisk) gives rise first to two synaptic boutons terminating in their respective dendritic tributaries (green and purple colored). Then, one of them (double arrowhead) gives rise to a descending (arrows) golf club like (GCL) running towards the capillary wall to end into a discreet enlargement (hollow arrow) next to the external basal lamina (not shown). Notice that both the dendritic and axonal components of this complex lie wrapped by the lateral aspects of the two end feet. F. Electron micrograph from a section of the series reconstructed in successive panels (i.e., G to J). The end-foot (blue) resting in the outer basal lamina (red) embeds partially and completely a dendrite (green) and various golf club-like expansions (assorted colors), respectively. The parent synaptic terminal (double arrow heads) contacts the dendritic spine (green) and gives rise to the GCL (arrows) anchoring next to the outer basal lamina (red) and ensuing longitudinal crest (hollow arrow). e = endothelium, PC = pericyte. G. Outer aspect of the complex basal lamina-longitudinal crest facing the overlaying end-foot, as seen in “F” or “I”. H. The synaptic bouton targets a pericapillary dendrite (green) and extends a GCL (both in transparent blue) close to the LC. I. Visualizing the end-foot (deep blue) illustrates its close interaction embedding the neural and interstitial (i.e., basal lamina) elements of the complex. J. Reconstruction of the five GCL extensions identified in the four microns encompassing the entire series. Notice that four of them lie close to the LC . Asterisk = parent axon; arrow = niche occupied by the spine shown in “F”. Scaling bars = 0.5 µm in A, D-J; 0.3 µm in C; 0.2 µm in B

**Figure 16.**
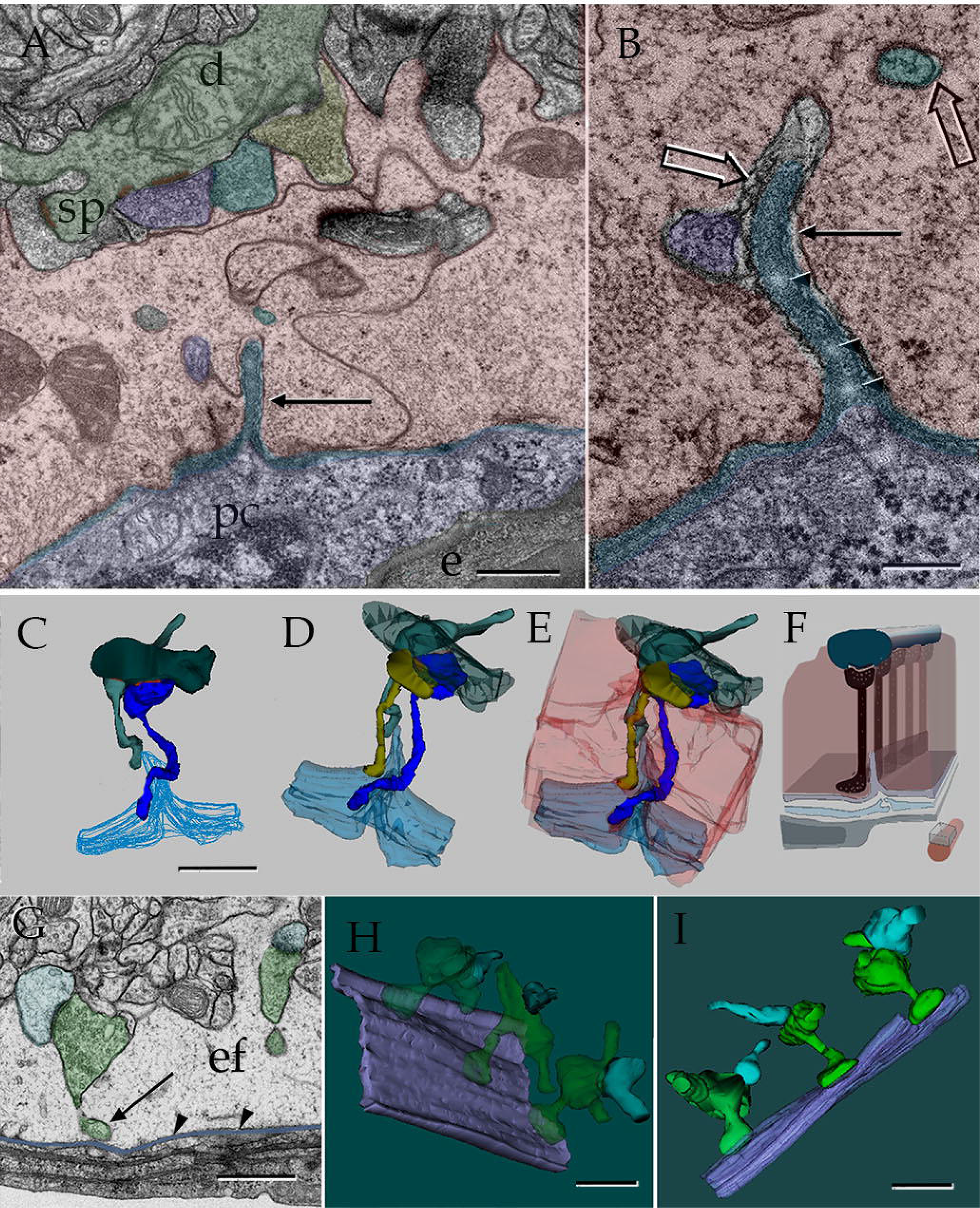
Series through a brain pericapillary unit (PCU). A. a longitudinal section through a dendritic shaft (d) bordered by several synaptic terminals. The end-foot (EF)(pink-colored) between the neuropil and external capillary basal lamina (oBL)(blue) arbors its longitudinal crest (LC) (arrow) as well as shafts of golf club-like (GCL) processes (purple and turquois) and their parent boutons synapsing the dendrite. e =endothelium; pc = pericyte. B. High magnification view of the end foot-end foot intersection organizing a distinct gap junction between them. Arrow heads = glycogen particles. C, D, E are progressive 3D views of the PCU. Two axo-dendritic synaptic terminals forming GCLs interacting with the LC of the oBL (outlined in blue). Dark green = dendritic shaft. D Three GCL bearing terminals converging about the LC. E. The neural elements as well as the basal lamina are engulfed by the EF which, together with the underlying endothelium-pericyte furnish the BPCU proper. F. Diagram of the PUC to whom the endothelial cell (gray) and pericyte processes (pale green) have been added. Notice that GCLs accommodate next to the LC along the longitudinal axis i.e., parallel arrangement. Scale bars 0.5 µm in A, 0.2 µm in B. G. An electron micrograph from the series reconstructed in “H” and “I”. Oblique sections through the capillary wall showing two axo-spinous synapses (green) originate two golf club-like extensions (GCL); in one of them (left) the terminal enlargement (arrow) lies next to the basal lamina (purple). Note the end foot (ef) surrounding synaptic boutons. Arrowheads = external basal lamina; light turquoise = dendritic spine. H and I are oblique and horizontal 3D views of the series encompassing three perivascular synaptic complexes with their respective GCL extensions aligned along the longitudinal axis of the capillary blood vessel. Scale bars = 0.5µm in A, 0.2 µm in B, 1.0 µm in C-E, and 0.5 µm in G-I

**Figure 17.**
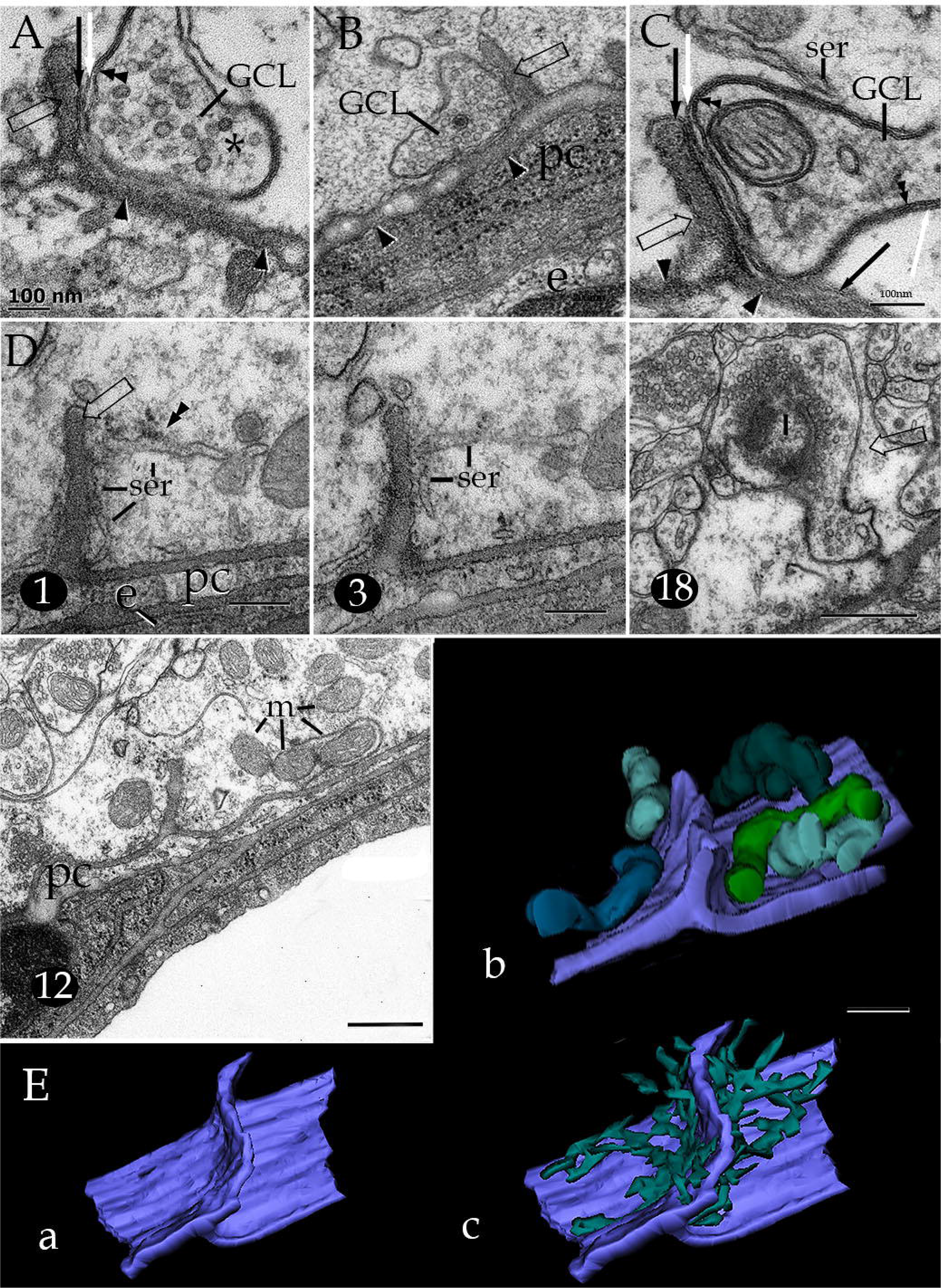
Electron microscopic views of golf club like synaptic extensions (GCL) and the neighboring pericapillary region. A. High magnification micrograph encompassing the interaction of a GCL next to a longitudinal ridge (hollow arrow). Between them, there are two fold interposing inner (white arrow) and outer (black arrow) plasma membranes of the end-foot. Note the numerous small clear, rounded vesicles contained by the GCL cytoplasm (asterisk). Double arrowhead = plasma membrane of the GCL; single arrowheads = outer capillary basal lamina. B. A GCL extension laying between the outer capillary basal lamina (arrowheads) and an arising longitudinal ridge (hollow arrow). Notice that the former contains numerous small rounded synaptic vesicles surrounding a solitary dense cored vesicle. pc = pericyte, e = endothelial cell. C. High magnification micrograph to depict the astrocytic, double plasma membrane (arrows) dissecting the longitudinal ridge (hollow arrow) and the GCL plasma membrane (double arrowhead); ser = smooth endoplasmic reticulum. D. assorted micrographs from a series encompassing 67 sections, position in the series is designed by Arabic numerals at the lower left of the figure. Organelles of the end-foot cytoplasm (ef) associated with the longitudinal crest (hollow arrows), consisting of smooth endoplasmic reticulum (ser) with scarce rough endoplasmic reticulum (double arrowheads). In section 18, the enlarged, terminal portion of two golf club like extensions (asterisk) next to the longitudinal crest are observed. m = mitochondria. E. The longitudinal crest and associated organelles are reconstructed from the series partly shown in “D”. a = external basal lamina and arising longitudinal crest b. mitochondria. c. smooth endoplasmic reticulum. Scale bars = 100nm A-C, 0.2 µm in D, 0.5 µm in section 12 and E.

**Figure 18.**
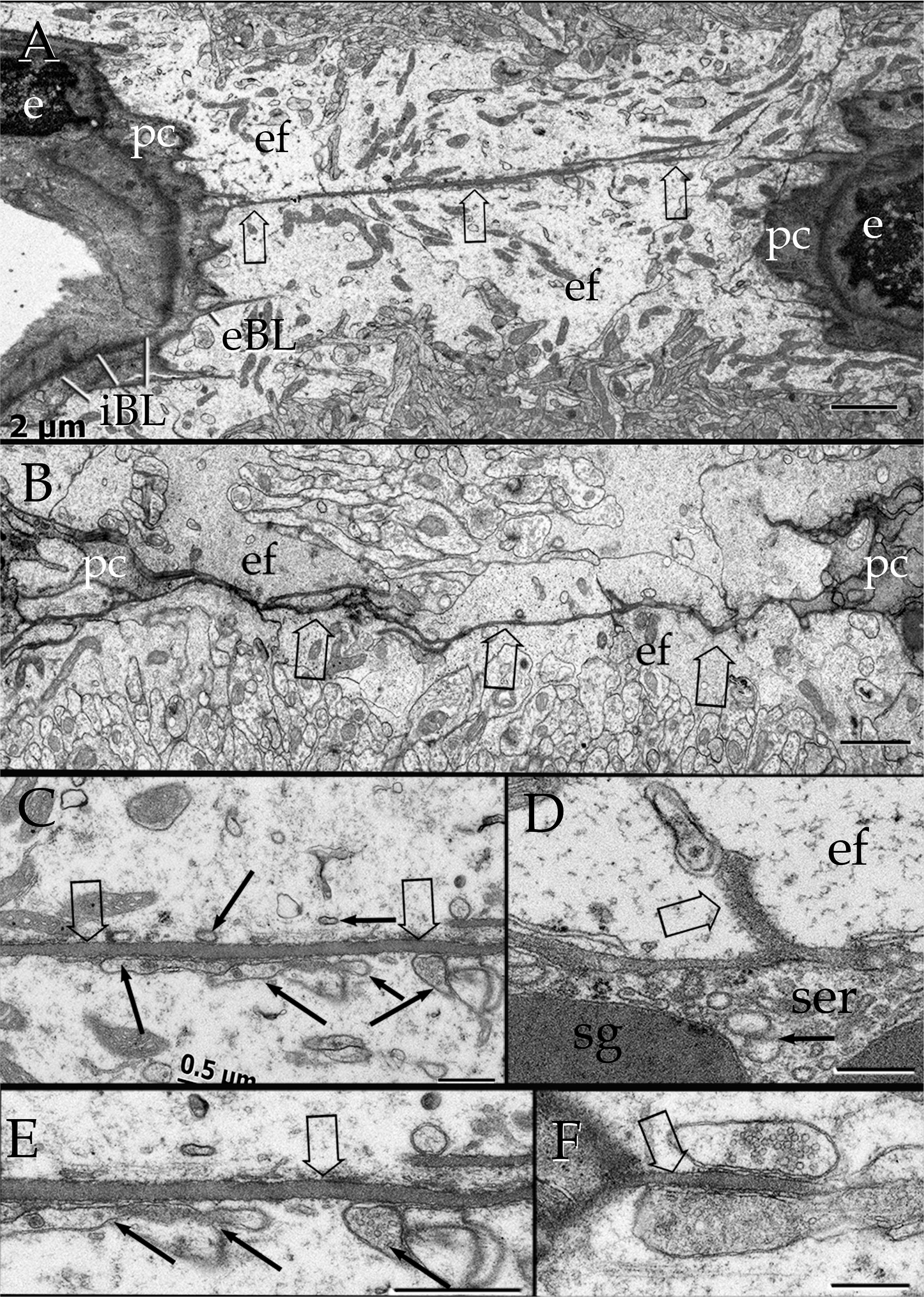
Assorted electron micrographs depicting horizontal sections through longitudinal crests (LC)(hollow arrows). A. The section discloses a LC between two transversely sectioned capillary blood vessels whose endothelial cells (e) and surrounding pericyte cytoplasm (pc) can be seen on either side of the micrograph. Notice the complete end feet (ef) covering of both capillary external basal laminae (oBL) and LC proper. iBL = internal basal lamina. B. Another oBL LC (hollow arrows) bridging two capillary blood vessels with pericyte processes (pc) associated with the capillary wall in either side (not shown). C. A segment of an oBL LC bounded by several terminal expansions (black arrows) of golf club-like processes. D. Transverse section through a longitudinal crest the oBL overlapping the cytoplasm of a Mato cell with large secretory granules (sg) surrounded numerous cisterns of the smooth endoplasmic reticulum (ser). E. High magnification view of that field outlined in “C”. Notice the thin cytoplasmic layer (arrow heads) of the end foot interposed between the LC and enlargements of the GCL. F. A LC bounded by two terminal dilations of GCLs. Note the pleomorphic vesicles and the higher electron density of the GCL at the bottom, contrasting with the rounded vesicles of the upper GCL. Scale bars = 2 µm in A and B; 0.5 µm in C to F.

Golf club-like processes (GCL). Among the broad category of pericapillary nerves (PCNs), GCLs are the most frequent ones in thalamic and neocortical specimens. A typical GCL arises from a committed synaptic contact that otherwise matches in every structural respect axo-dendritic chemical synapse (Peters et al., 1976), defining a modified tripartite synapse (Araque et al, 1999)(mTS). Unlike typical tripartite synapses, surrounded by a peripheral astrocytic processes, 3D reconstructions of pericapillary mTS, usually axo-dendritic, are surrounded by the EF proper (Varela-Echavarría, 2017). The minute and variable dimensions of a GCL (i.e., 0.3 to 5µm in length and 0.2 µm diameter) require of reconstructions of series to be fully visualized. A typical GCL consists of funnel-like out-growth descending for a variable distance in direction to the oBL (see above), opening to the enlarged, end-sac part of the GCL (Figs. 15E, F, H, J; 16G-I, 17A-C). Vesicular contents of the parent bouton and the arising GCL include numerous small, agranular vesicles that, together with a dendritic spine or shaft (15E, H, 16A), define an asymmetrical synaptic contact (Peters, et al, 1976). Both the stem and the end sac of the GCL contain scattered rounded synaptic vesicles (Fig. 17 A, B) with scarce rounded electron dense core vesicles (Fig. 17B). GCLs containing flat, pleomorphic vesicles are uncommon (Fig. 18F), averaging two for every fifty GCLs. Careful observation to the GCL cytoplasm fails to identify an active zone(s) and/or junctional complexes other than that of the parent bouton. While a neighboring, putative effector structure of the GCL notwithstanding, clusters of synaptic vesicles appear to be polarized toward the oBL.(17B, D). Because of the above-mentioned GCL engulfment by the EF, the interposed inner and outer cell membranes lie squeezed between the former and the oBL (Fig 17A-C). GCLs are ubiquitous throughout central CBVs and we have additionally identified them in series from the olfactory bulb and peduncle (not shown). The outstanding GCL structure is commonly reciprocated by the interstitial ground substance as commented upon next.

Interstitial ground substance. To the oBL specializations described for the PC-AP, there is still another heretofore unnoted structure in CBV throughout the forebrain. Transverse sections through the CBVs disclose two to five small outgrowths of the abluminal aspect of the oBL, reciprocating with the longitudinal concavity just described for the EF (Fig. 15D). This sort of outgrowth, termed here longitudinal crest (LC), approximates 0.5 to 2.0 in height by 0.3 microns in width (Figs. 16 A-F, 17). Outstanding, sections paralleling the CBV longitudinal axis, reveal that the LC extends along the entire CBV length (Fig. 18).

Although the LC is an homogeneous extension of the oBL matrix, clusters of one to four collagen fibers may be seen embedded in about a third of them (Figs. 16A,B). Shape and dimensions of the LC are uniform throughout the brain CBVs but are variable in the cerebellar cortex (see below). The LC association with the terminal expansion of the GCL appears to be a fundamental site for neurovascular interaction (Figs. 16A-F, 17B-D). Direct interactions of the GCL with areas of the oBL devoid of LC were uncommon in samples from the thalamus and isocortex (Figure 16 G-I).

At stage of our description, it is advisable to turn our attention back to panel Figure 16. The dendritic shaft (d) of a thalamic neuron receives a series of consecutive boutons sending each a GCL that anchor in an identical, succession along the LC, as defined by reconstruction of the series. The peculiar sequence of synaptic bouton-GCL-LC also observed in the cerebral isocortex (Fig 18C), and basal ganglia (not shown) is suggestive of ubiquity. Figure 19 summarizes these descriptions and shows the neuro-glial capillary unit (NCU) as a whole. The arising question about the possible source(s) of prospective pericapillary dendrites receiving synaptic-bouton-GCL-LC is approached subsequently.

**Figure 19.**
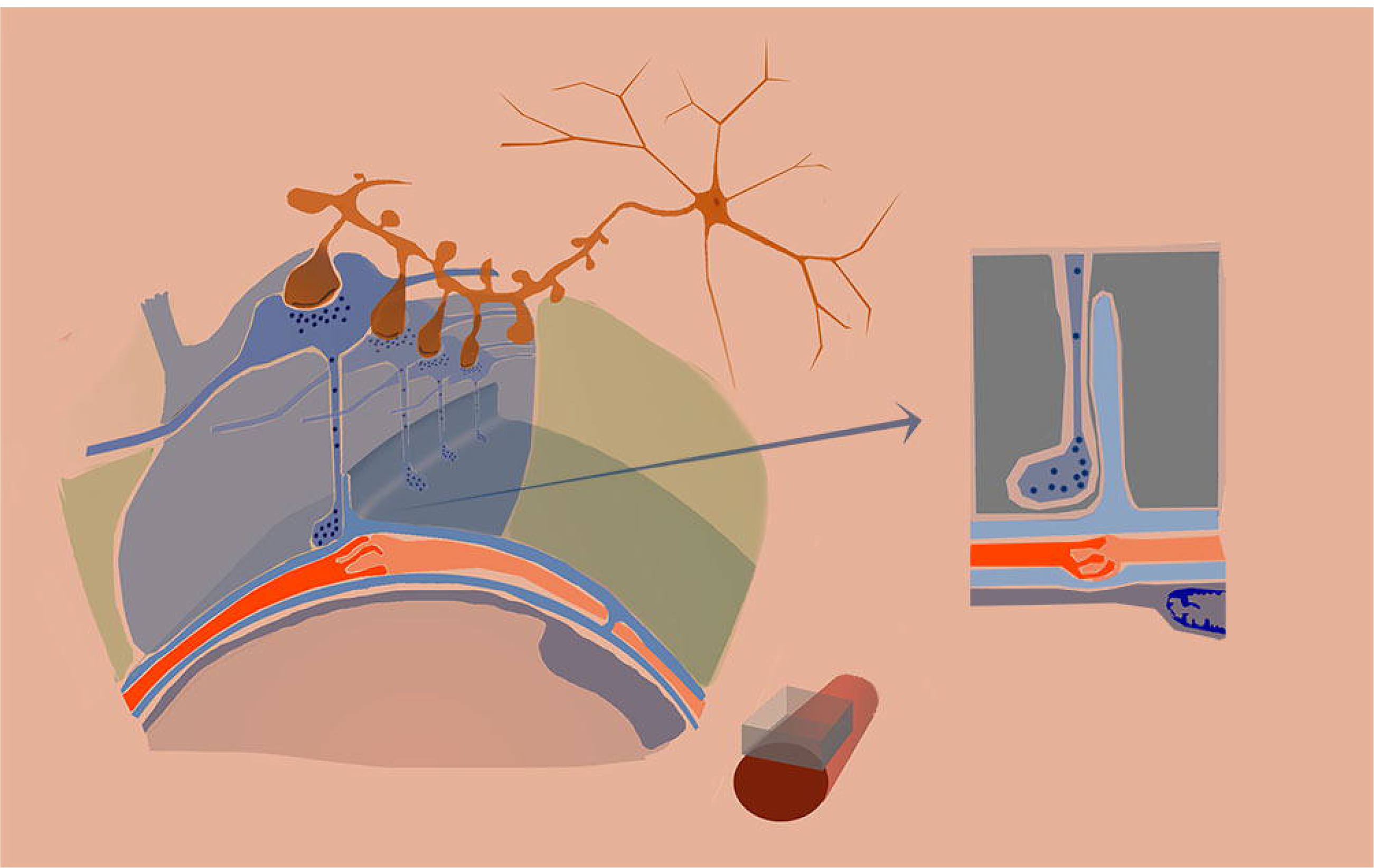
Diagrammatic representation of the neuro-glial capillary unit (NCU). The endothelium and interlacing pericytes (PCs) (orange) are surrounded by the inner- and outer basal lamina (BL) and the former interacts with the blood flow (light gray). Underneath the PCs structure a cellular continuum interlacing with successive and adjacent capillaries. The specialized longidudinal crest (LC) protrudes from the outer basal lamina opposing to the terminal portions of golf club like extensions (GCL) resulting from the descending outgrowth of the axo-spinous dendrite. Both the dendrite and the GCL-synapse are surrounded by the EF which prevents the direct interaction between GCL and LC (right side) (gray). A tributary dendrite receiving the orderly synapse-GCL-LC series together with the EF covering, and the PC-capillary wall define the NCU.

A set of distal dendrites from cortical pyramidal (Fig. 20), medium size spiny (Somogyi and Smith, 1979), thalamic projecting (Clasca et al., 2012) cerebellar-(vide infra) granule cells (vide infra) undergoes structural modifications at their respective vascular intersections. These pericapillary changes may be summarized as first, contrasting with the cylindrical or varicose appearance of the parent dendrite is the occurrence one to four balloon-like enlargements of the parent dendritic shaft resulting in a tuberose appearance (Fig 21A B, 22E). These rounded outgrowths exhibit, in turn, assorted membranous appendages of variable length. A second dendritic specialization observed in distal dendrites of pyramidal neurons (20A, 21D) consists of short and narrow, supernumerary branches, covered by a few pleiomorphic, atypical spines (Fig 21D) . Farther distally the tributary dendrite(s) resume its former structure. While most dendritic-CBV interactions take place of passage few terminal branches end-up in the perivascular domain (Fig 21C, 22E). Still another signed feature of the dendrite-CBV interaction is that the translucent EF mantle surrounding them (Figs 20A, 21A). Since, pericapillary dendritic interactions of the NCU represent an obvious candidate for the GCL-synaptic bouton output a search for their respective parent neuron and axonal tributaries is performed next.

**Figure 20.**
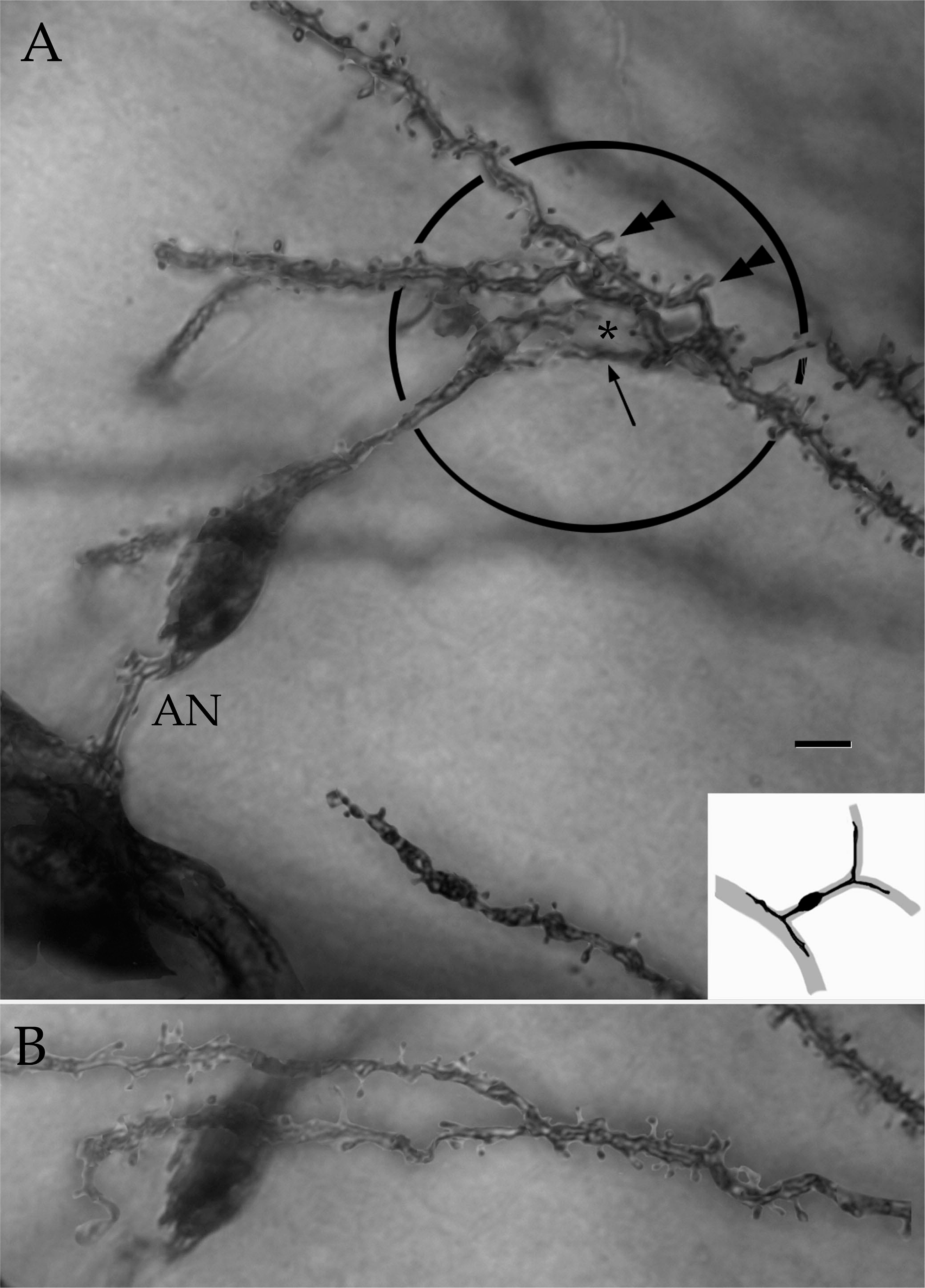
Light microscopic digital photomontages of dendritic and axonal processes intersecting with the perivascular neuropil. A and B. Distal pyramidal cell dendrites anchoring next to the secondary processes (circle) and perikaryon (pc) of a nearby pericyte (black-colored in inset; d = dendrite). Notice that the overlaying dendrites originating thicker spines (double arrowheads) and thin supernumerary branchlets (arrows). Notice the glial mantle (asterisk) surrounding dendrites and distal pericyte processes. AN = axonoid with a discreet flexure. B. Two perisomatic dendrites anchoring the cell body of the pericyte shown in “A”. C. Perikaryon of the pericyte in “A” with two opposing pyramidal cell dendrites (arrows). Scale bars = 1µm

**Figure 21.**
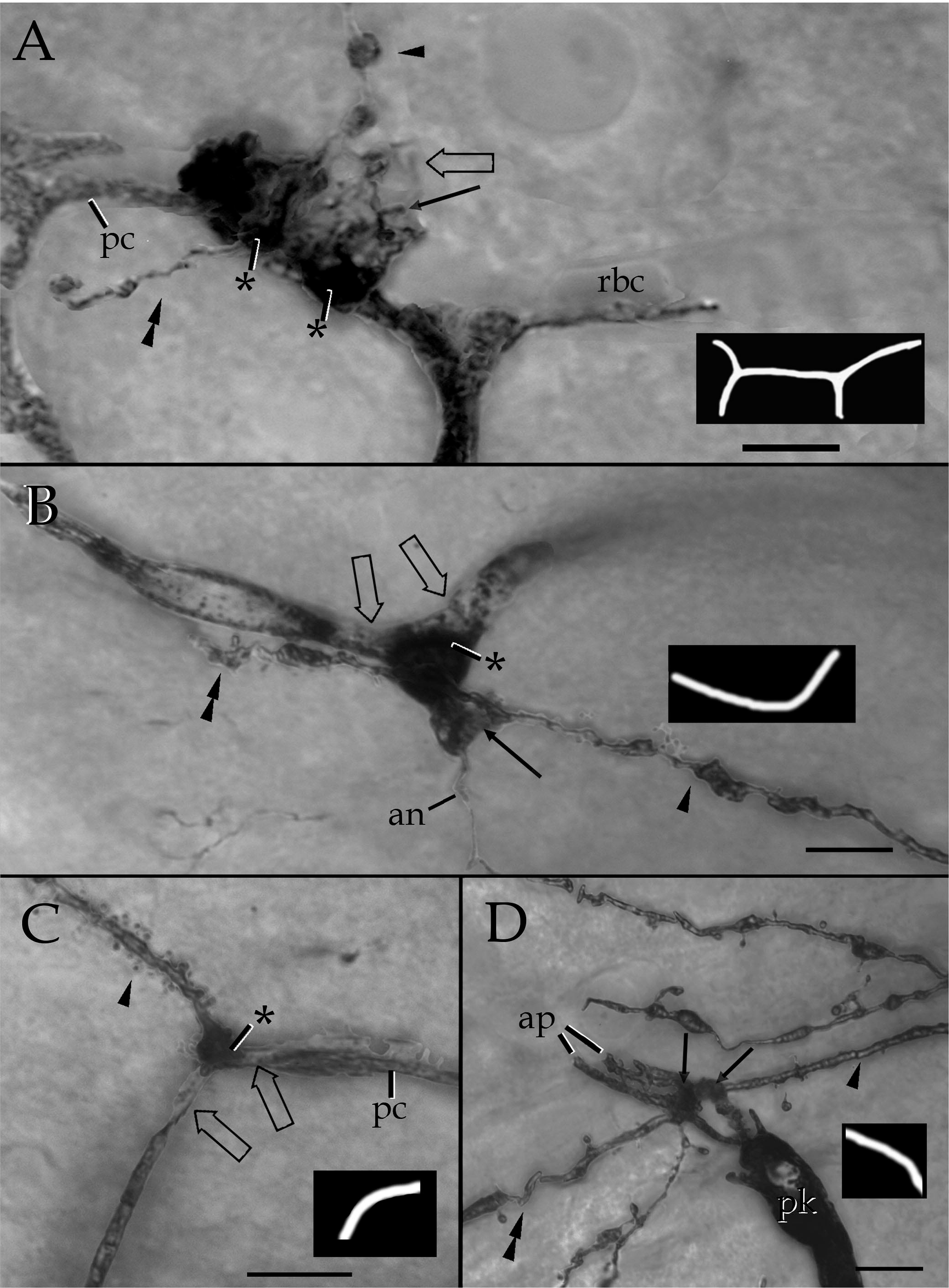
Structural variations of dendritic processes within the pericapillary domain. The diagram at the lower left defines the capillary trajectory. A. A terminal dendrite of an interneuron coursing from proximal (arrowhead) to distal (double arrowhead) within the pericapillary area assorted out growths (arrow) both of which appear masked by a glial envelope (hollow arrow). pc = pericyte processes; rbc = row of unstained red blood cells contained by the capillary blood vessel. Parietal isocortex. B. A solitary granule cell dendrite extending from proximal (arrowhead) to distal (double arrowhead). As the dendrite reaches the capillary wall it originates a large, rounded enlargement (asterisk) engulfing the pericyte process (hollow arrows) surrounding the blood vessel. an = perforating axonoid . Terminal dendrite (arrow head) of a medium-sized spiny cell that end abruptly into a single, rounded enlargement (asterisk) bounded by two pericyte processes (hollow arrows). Caudo-putamen nuclei. D. Terminal dendrites of thalamic neurons, one of them overlaps to the transverse limbs of the asymmetrical processes (ap). pk = perikaryon of a pericyte. Lateral thalamus. Scale bars = 3µm.

The previous descriptions dealt with short interactions of perivascular nerve processes with glial and intercellular specializations associated with the CBV. However, is insufficient to reveal the neuron-types providing extensions the neuropil-CBV domain. A search for neuronal interactions with the CBVs in the cerebellar cortex is performed by the combination of the Golgi technique with transmission electron microscopy and selected 3D reconstructions.

Contributing neurons to the pericapillary neuropil. Inspection of the granule cells in the homonymous layer (GC) layer of the cerebellum reveals various neuro-vascular specializations (Figs. 22 and 23). A systematic search revels that occasional granule cells dendrites entangle the capillary wall. For instance, the GC dendrites originate globular structures stung by fine cytoplasmic bridges in intimate association with the capillary wall (23A). A dramatic interaction of this dendritic-CBV association occurs when a side branch from the dendritic shaft extends a thin, long collateral of large spherules stung by thin bridges that entrap the capillary wall for a variable distance (Fig 22E). Although, the GC axon ascends unbranched to the molecular layer (Ramon y Cajal 1904, Palay and Chan-Palay 1974) our observations depict short and long axon collaterals to the CBV. Perhaps the most dramatic, consists of a very short collateral made up of numerous rows of anastomotic vesicles masking the PC processes (Fig. 22A, B) Electron microscopic observation of dendrites at the GC-CBV interaction uncover that some globular structures arising from proximal dendrites are indeed small oval extensions facing the oBL (Fig. 23A). 3D ensembles further visualize that the surrounding EF provides an uninterrupted envelope to the GC outgrowths (Fig. 23C). While appositions of the GC perikaryon itself to the CVB are uncommon, occasional finger-like outgrowths, adjacent to the CBV wall are observed encased by the EF (Fig. 23D)

**Figure 22.**
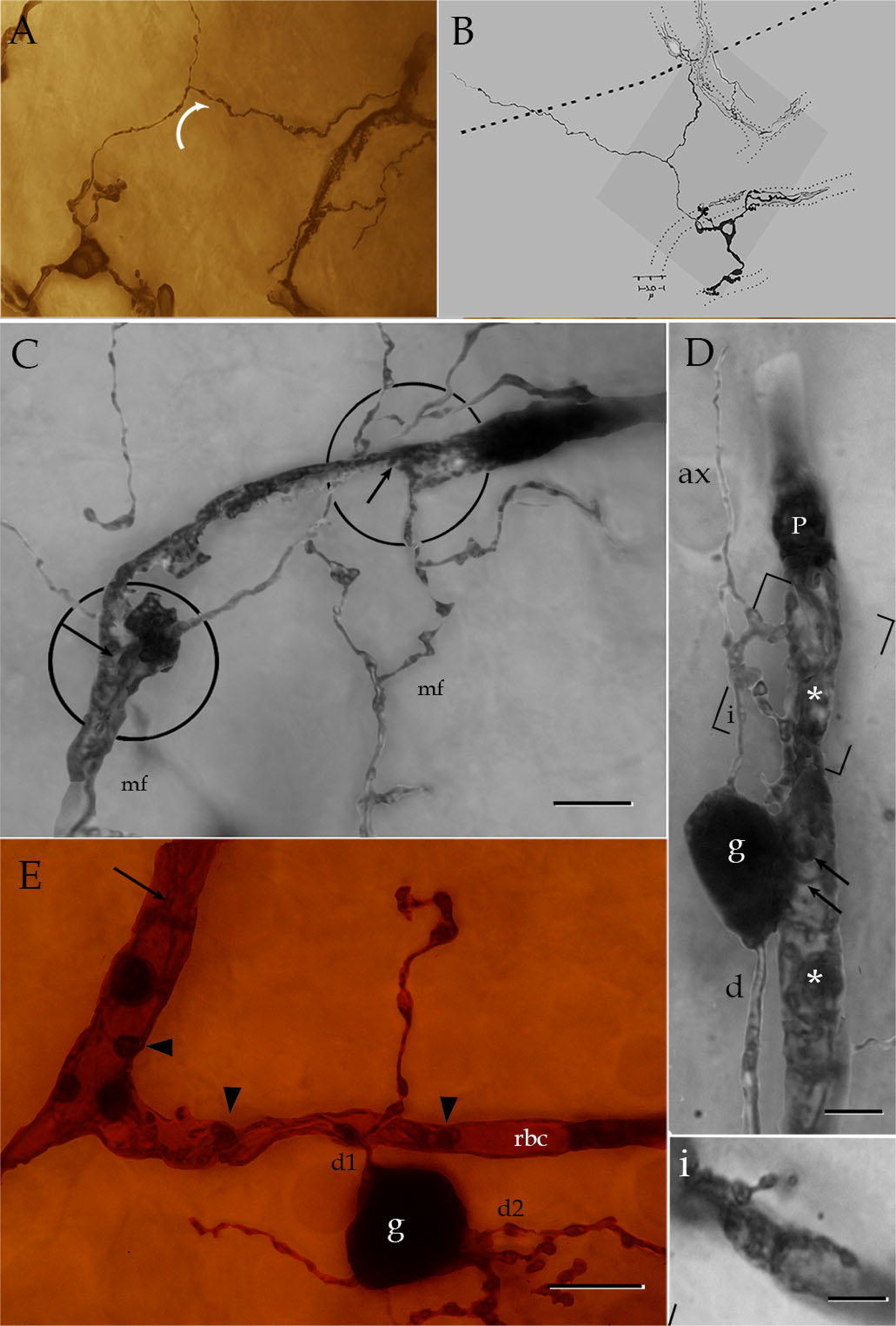
Light microscopic views from the granule cell layer of the cerebellar cortex. A. Photomontage from a granule cell with three divergent dendrites. In its ascending path the axon sends a thicker collateral (arrow) that entangles with a capillary blood vessel. B. Camera lucida drawing of the neuron shown in “A”. The lower magnification of the field shows the interaction of both the granule cell axon collateral (upper right) and dendrites entangling the capillary wall (dotted). Horizontal dotted line defines the boundaries between the granule cell- and molecular layer. C. Photomontage of the granule cell layer. Two ascending mossy fibers (mf) interact with pericytes by means of a mossy terminal (left) or short tongue-like excrescences (arrows). See Fig. 22D. D. Photomontage of the intersection of the granule cell (g) perikaryon and its proximal processes with a pericyte surrounding the capillary blood vessel. Notice the massive ramification of the axon collateral encircling (asterisks) the underlying pericyte (p) as well as the short outgrowths of the perikaryon (arrows) (see Fig 21 D). ax = distal axon, d = dendrite. i. Single electron micrograph from the photomontage shown in “D”. Notice the ramifying granule cell axon collateral encircling the external aspect of the capillary wall. E. Granule cell originating two divergent dendrites (d1 and d2), one of them (d1) ramifies next to the cell body into an horizontal branch embedded in the capillary wall (arrow heads). Notice the distal processes (arrow) from a putative pericyte. rbc = red blood cell. Scale bars = 5 µm in C and E, 2 µm in D, 1 µm in i.. Adult rabbit, rapid Golgi technique.

**Figure 23.**
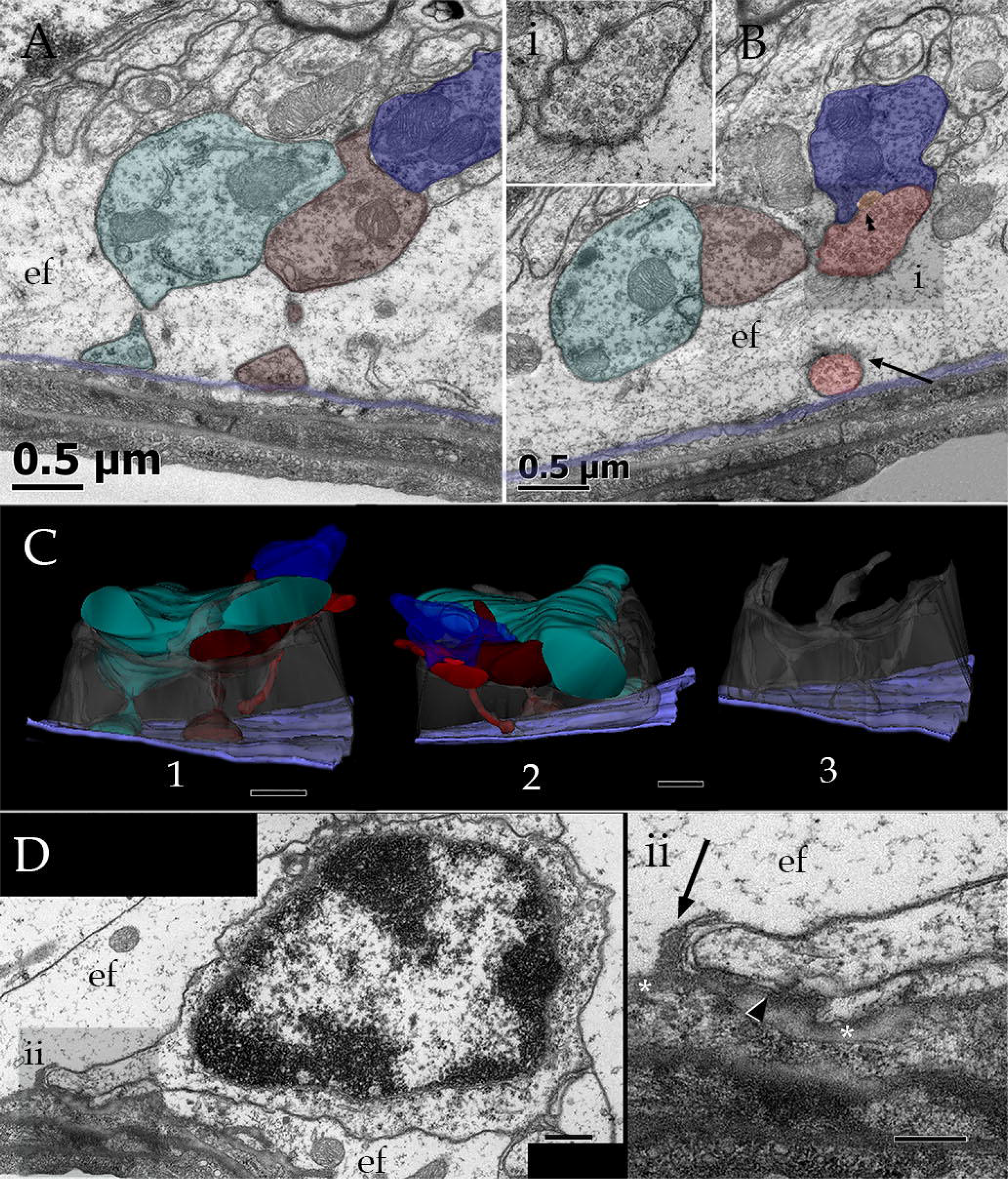
Electron microscopic views of the granule cell layer of the cerebellar cortex. A. paired granule cell dendrites (green and pink-colored) originating homonymous spines laying next to the outer capillary basal lamina (arrow). B Complementary micrograph from the same series than that of “A” in which an additional dendrite (blue) is contacted by a putative synaptic terminal from an stellate cell whose flat vesicle-containing bouton (red) gives rise to a golf club like extension (arrow). Note that the latter anchors next to the outer capillary basal lamina (light purple). Arrow heads = synaptic active zone. i = high magnification view depicting the flat-type of vesicles contained by the synaptic bouton. C. Reconstructs from the complete series. 1 and 2. include dendrites (blue, green and deep red). In 2 a Golgi cell axon whose synaptic terminal extends a golf club like (bright red) contacting a dendritic shaft (blue) and descending next the basal lamina (light purple). Bright orange = active synaptic zone. 3. Image with exclusion of the nerve processes visualizing the glial envelope provided by the end-foot (gray). precluding direct contacts between the former and the basal lamina. Orange = active synaptic zone. D Electron micrographs from a granule cell soma originating a perivascular pseudopod anchoring next to the outer capillary basal lamina. Notice that the cell is completely engulfed by the end feet (ef). ii. High magnification view disclosing the interaction of the granule cell pseudopod with the basal lamina (asterisk) and an arising longitudinal crest (arrow), between them and the former a thin cytoplasmic sheath (arrow head) precludes direct contact. Calibration bars = 0.5 µm in A to D, 0.2 µm in insets.

Mossy fiber (MF) represents the best-studied afferent fiber-system of the cerebellar cortex. The ascending MF is a branching myelinated axon that alternates with a large synaptic complex, or rosettes whose appearance confers the name to this incoming fiber (Ramón y Cajal.1904, Chan and Palay 1971; Hamory and Somogyi 1983; Jakab and Hámori 1988). Our observations of the GC layer add the presence of peculiar MF-CBV intersections (Fig 24A). Directed search for these interactions exposes a distinct rounded to oval extrusions of the rosette next to the PC processes (Fig 24B). These outgrowths are pleomorphic but most of them drumsticks (Fig. 25B2,3) or small swellings. CBVs may occasionally be engulf by a rosette (Fig. 24B1, 25A). Reconstructions of MF at the superficial part of the granule cell layer (Fig. 26) complement light microscopic observations showing that when the MF approaches the CBV, it extends an outgrowth (vide supra) and both acquires an EF envelope (Fig. 26).

**Figure 24.**
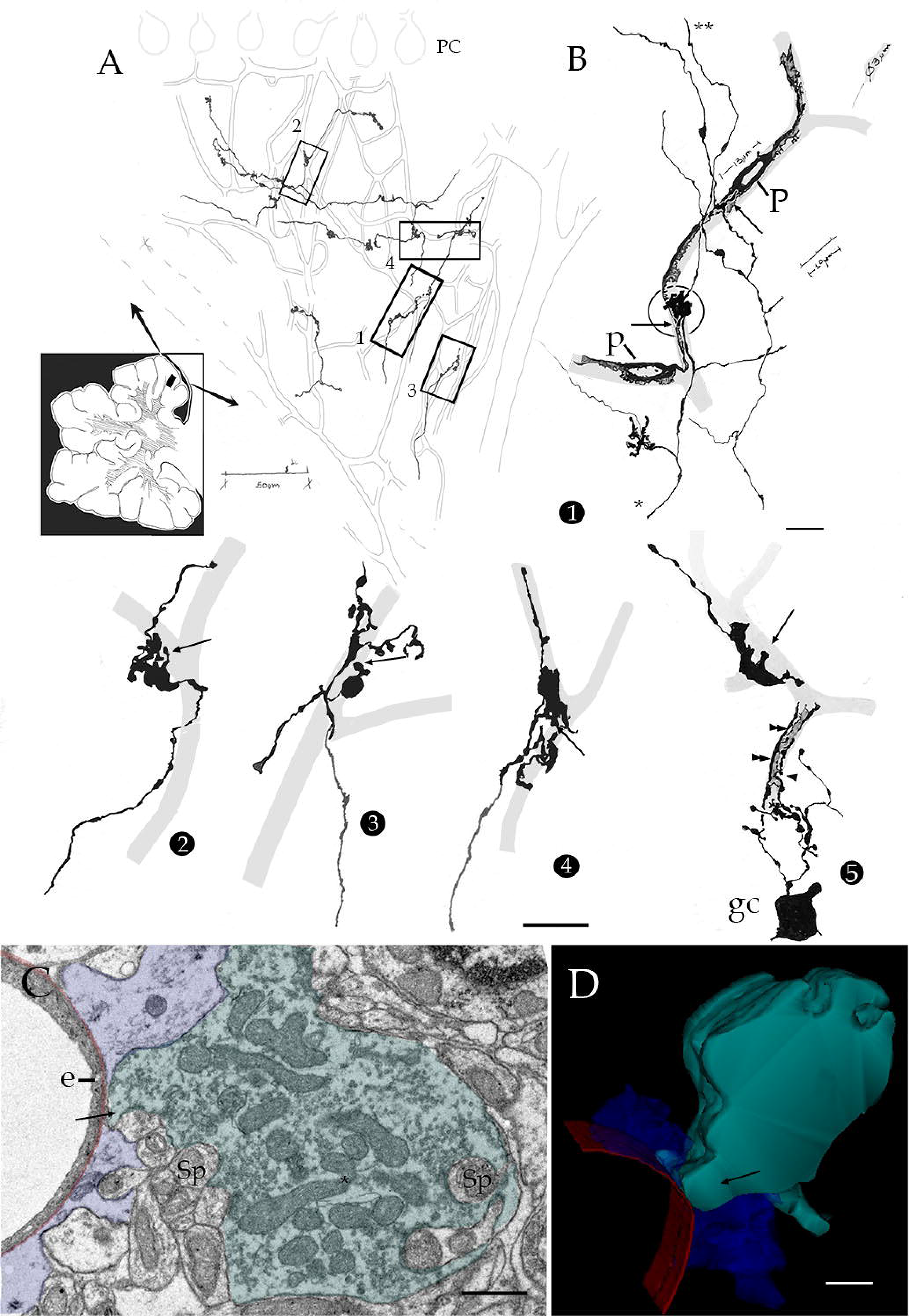
Light and electron microscopic views of mossy fibers in cerebellar cortex . A. Camera lucida drawings of blood vessels in the granule cell layer (soft pencil). Mossy fibers whose synaptic boutons overlap capillary blood vessels have been drawn with black ink. B High magnification drawings from mossy fibers and synaptic boutons. Drawings designated 1 to 4 were obtained from that section shown in “A”. 1. Two ascending mossy fibers whose boutons are interact with pericytes (p). Notice that the synaptic boutons extend short of passage outgrowths (arrows). Capillary blood vessels are drawn with soft pencil. asterisk = proximal mossy fiber, double asterisk = distal part of the mossy fiber. 2, 3, 4 and upper fiber in 5 are of passage outgrowths (arrow) from mossy fiber boutons. 5. A dendritic collateral from the granule cell (gc) lies in apposition to the secondary process (double arrow heads) of a pericyte (not shown). C. A pericapillary mossy fiber terminal (green) protruding to the capillary basal lamina (red), bounded by an end foot (blue). e = endothelial cell, sp = granule cell dendritic spine. D 3D view from the mossy fiber terminal whose distinct outgrowth (arrow) penetrates between the end foot (blue), next to the basal lamina (red). Notice the indentations at the upper right of the bouton corresponding to granule cell dendritic spines. Calibration bars = 50 µm in A, 10 in B, 0.5 in C and D.

**Figure 25.**
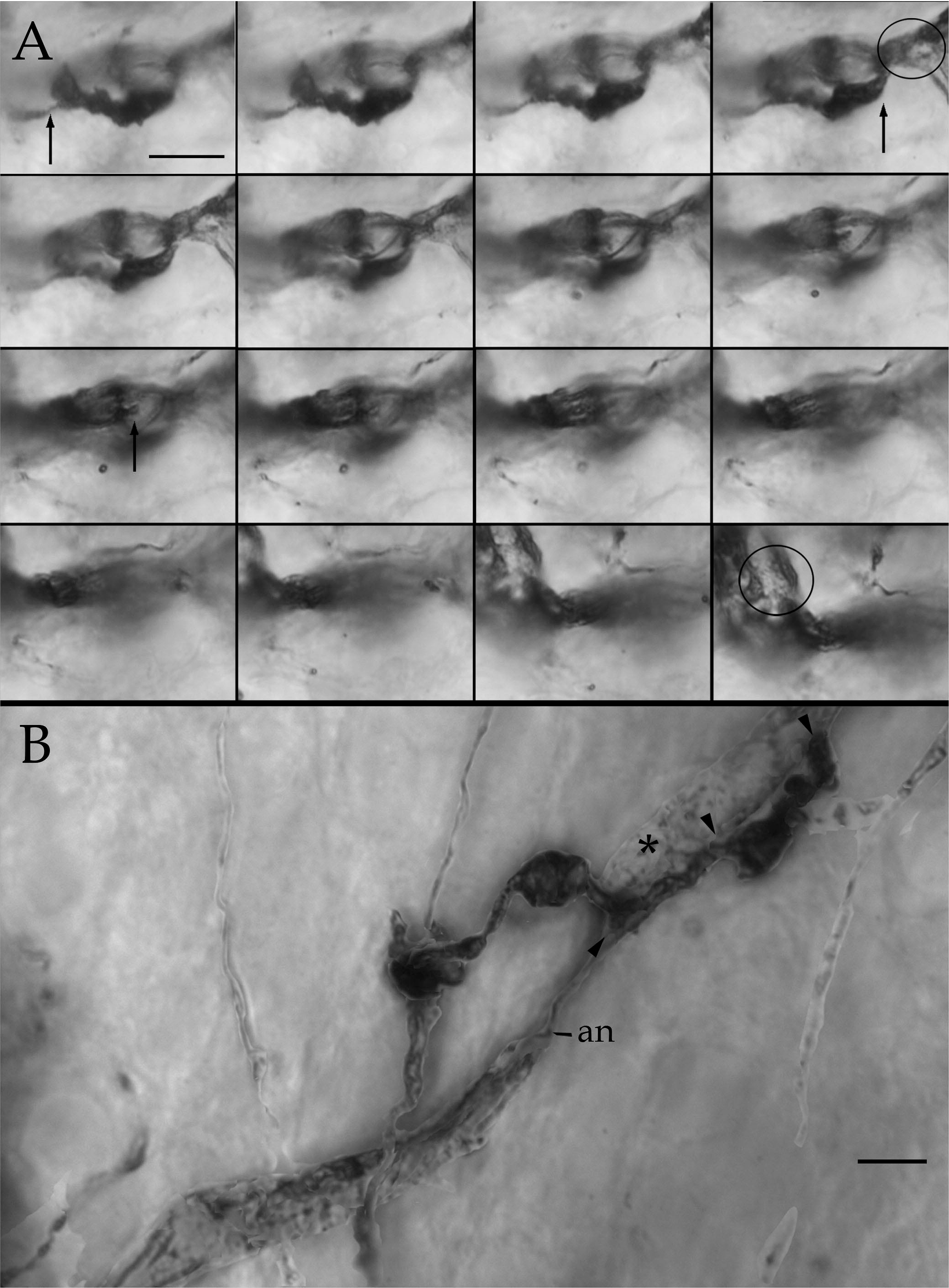
Light microscopic photomontages of mossy fibers. A. consecutive pictures at different focal depth illustrating the superposition of three synaptic outworths to the capillary wall. Notice the pericyte processes (circle) alternating with the synaptic outgrowths (arrows). B. A singe mossy fiber alternating with three rosettes. Notice that the axonal shaft a three rosettes two of them extending short outgrowths overlapping the pericyte processes (asterisks). an = axonoid. Rapid Golgi stain. Calibration bars = 2 µm.

**Figure 26.**
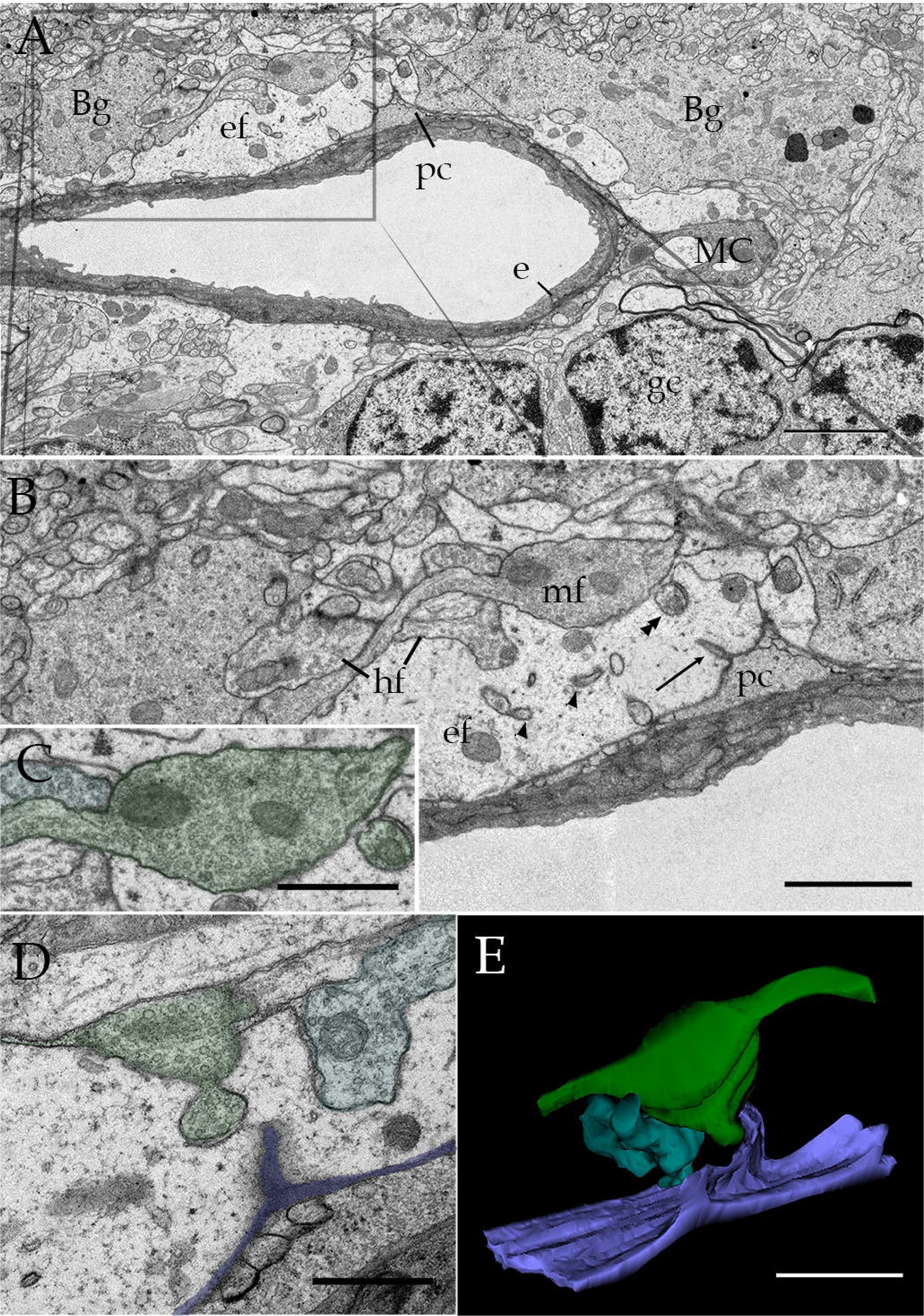
Electron micrographs of the pericapillary neuropil at the superficial aspect of the cerebellar granule cell layer. A. Tangential section through a capillary blood vessel surrounded by Bergmann glial processes (Bg), Mato-(MC), granule-cells (gc), end feet (ef) and nerve processes. B. High magnification view of that part of the area labeled in “A”. Longitudinal section through a horizontal-(hf) and mossy fiber (mf) axons. The capillary wall surrounded by pericyte process (pc) underneath a longitudinal crest (arrow), and electron-lucid end feet (ef). The later are pierced by penetrating fibrils (arrow heads), possibly golf club-like processes. C. High magnification view of the mossy-(center) and horizontal (upper left) fibers. Notice the tiny vesicle containing appendage from the former. C. micrograph for another section of the series in which a frank protrusion is budding of from the mossy fiber terminal (green).A longitudinal crest (blue) covering the pericyte process is seen. E. 3D image from the series shown in A to D. Notice the synaptic outgrowth of the mossy-(green) and horizontal (turquois) fiber terminals directed to the LC-basal lamina (purple). Calibration bars 2 µm in A, 0.5 µm in B to E

Ramón y Cajal (1904) described that blood vessels imprint in the Purkinje cell dendrites areas devoid of spines. These dendrites can be seen in our specimens (Fig. 27A). Electron microscopic observations at the Purkinje cell dendrite-CBV appositions reveal that a unique neuropil may lay squeezed therein. Contrasting with the radial arrangement observed throughout the dendritic tree of this cell type (Lee et al., 2005)It consists of sparse bent dendrites receiving axo-spinous synapses, some of them engulfed by the EF cytoplasm (Fig. 27B, C). The later synaptic terminals appear to correspond to those described for the parallel fiber bouton by Palay and Chan-Palay (1974). Further, the parallel fiber-Purkinje dendrite synapse-GCL is surrounded by an EF. Unlike most interactions of the GCL-LC observed in both cerebral cortex and thalamus, no LC is seen, although like elsewhere in the forebrain tight PC processes lie under the smooth oBL (Fig 26B, 27C). Purkinje cell dendritic spines receiving climbing fiber synapses may also be seen encased by EF (Fig. 27E-G). No 3D ensembles from these synapses are available.

**Figure 27.**
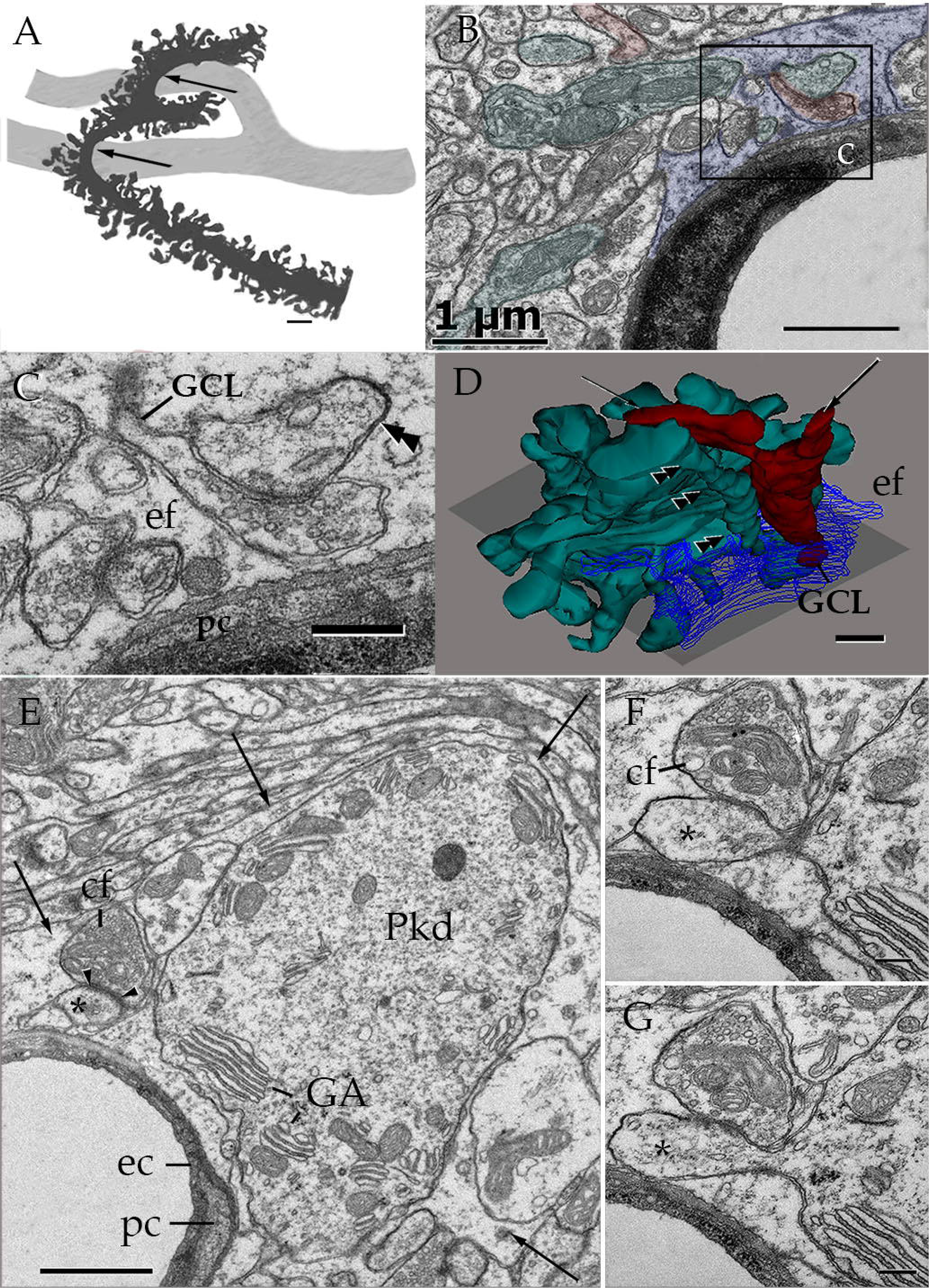
Purkinje cell dendritic spines interacting with parallel fiber terminals within the capillary wall. A. High magnification camera lucida drawing showing the impressions left (arrows) by two blood vessels (soft pencil), on a proximal dendrite of a Purkinje cell (black)(PC). B, C, and D, feature serial sections through a PC dendrite (green) Climbing fiber (red) gives-rise to a golf club like extension (GCL) whose stem had contacted the protruding dendritic spine (double arrow heads) stems of the climbing fiber (arrows). In D, the end foot (blue) has partially been reconstructed (ef) disclosing the full covering to both, dendritic and GCL processes. Note in C the numerous small round, clear vesicles contained by the synaptic bouton originating the GCL. E to G are semi serial sections through a Purkinje dendritic process (PkD) extending a dendritic spine (asterisk) encased by an end-foot (arrow) in what appears to be a tripartite synapse. GA = Golgi apparatus. Calibration bars = 2 µm in A, 1 in B, D, and E, 0.2 in C, F, and G

Just above the row of Purkinje cells, the basket cell stands out for its unique axon, entangling their somata. In our sample, the two sets of axonal branches described earlier (Ramón y Cajal, 1904) are identified (Fig. 28A). One of them defines thick axon terminals about the Purkinje cell (i.e., baskets). Another sort of axons, usually beaded fibrils (Palay and Chan-Palay 1974), commonly arises from the parent thick axon ascending to the cerebellum surface. However, we found that baskets may in turn originate occasional beaded axons that ramify profusely encasing CBVs. (Fig. 28A; C). Serial sections of the intersection of granule cell-molecular-layer reveal that basket cell axons originating pericapillary terminal boutons. These boutons contain oval and flat synaptic vesicles and also lack a discernable target (Fig. 28D, E). Although EF fully encases these terminals too, no close apposition of them with the oBL may be identified.

**Figure 28.**
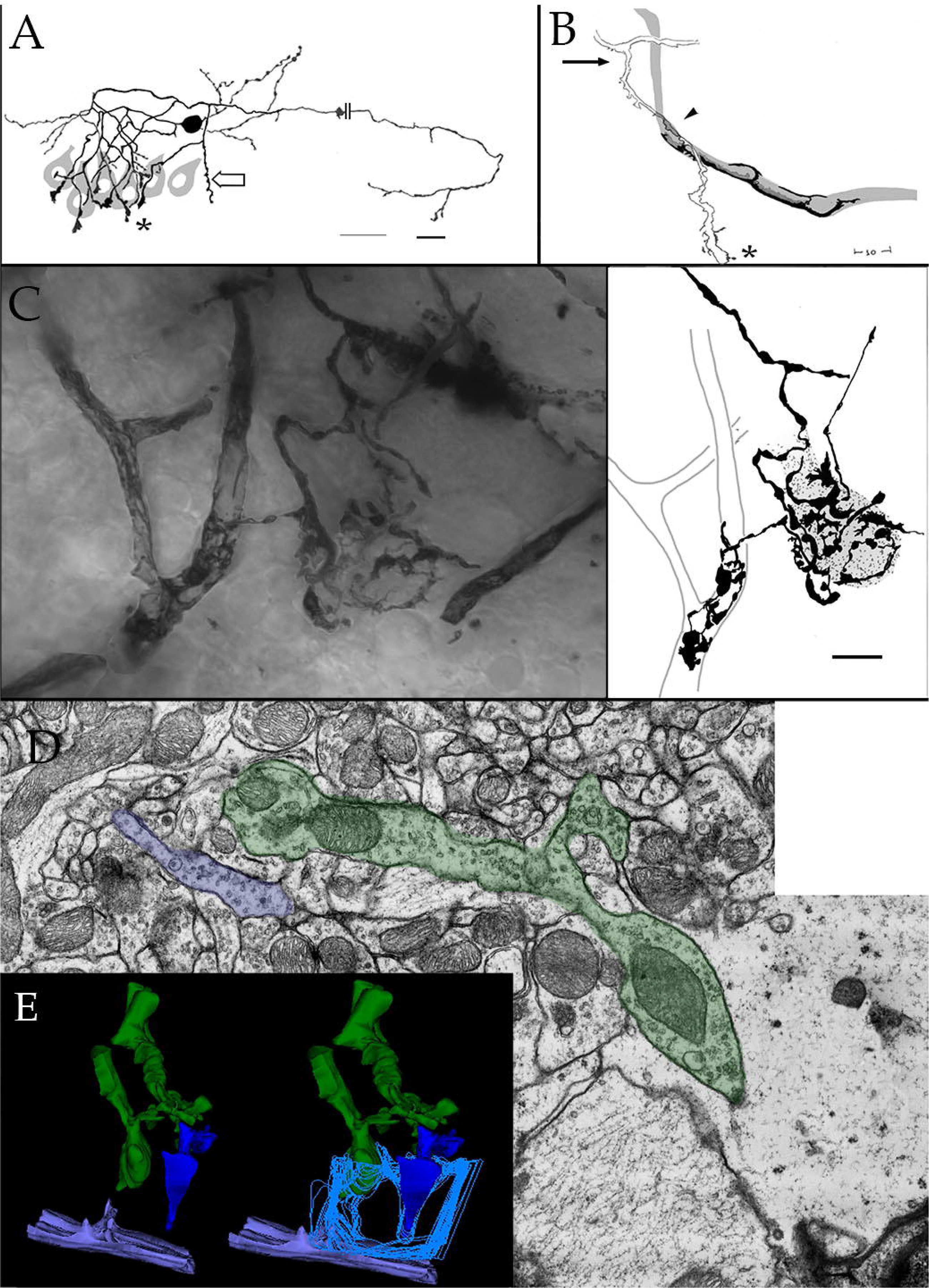
Light and electron microscopic views of cerebellar Basket cell and its perivascular axon processes. A survey camera lucida drawing of a basket cell in that part of the molecular layer intersecting with Purkinje cell somata (gray). The long axon of the former gives rise to thick, vertical extensions, or pinceau that insinuate about the later. Thinner, varicose fibrils (hollow arrows) are the second axonal specialization. B High magnification drawing of the root of a thick collateral (arrow) surrounding helicoidally a nearby capillary blood vessel (soft pencil) encased by pericyte processes (black). Asterisk = pinceau. C. High magnification views of a pericellular basket surrounding an unstained Purkinje cell perikaryon. Sidewise, the basket extends a varicose-type axon that, in turn, entangles the capillary wall, masking the pericyte processes (arrow). The full axons are drawn in the right side of the figure, outlying the capillary contour and the purkinje cell soma (soft pencil) D. Electron micrograph from a series illustrating parts of two presumptive distal axons (colored) of a basket cell. Axon endings (one not shown) (green) contain oval and, occasional flat synaptic vesicles and a single mitochondrion E. 3D reconstructs of the full ninety three sections. Left side reconstruct includes the two fibrils with their respective boutons (green and blue) and the capillary outer basal lamina with a protruding longitudinal crest. The end foot (light blue) enveloping both terminals is outlined at the right side reconstruct . Calibration bars in A to C = 10 µm, 0.5 µm in D, E..

### Blind nerve endings

While no putative nerve terminals are found in CBVs distributed in the diencephalon and forebrain, two distinct sorts of myelinated fibers appear to terminate in those of the cerebellar cortex. A first sort consists of 0.5 to 1.5 myelinated axons that as their stem axon approaches the CVB their axoplasm is filled-up by two or three mitochondria (Figs 29 A, C). Then, the axons lose their myelin envelope (Fig. 29B,C) and the axoplasm spreads into irregular endings which lacks synaptic vesicles with few coat vesicles. A second type of myelinated axon that appears to terminate in the capillary wall is a thinner ending approximating 0.75 micrometer in diameter. Unlike the previous one the axoplasm is virtually devoid of mitochondria and as soon as it approaches the capillary wall losses the myelin envelope. Like the axon itself the terminal complex contains few microtubules and its cell membrane underscored by a fussy material containing scattered coated vesicles (Fig 29 Fi). Like the mitochondria-load terminal, the distal portion of this axon subtype lies next to that part of the oBL covering the PC perikaryon or its processes. Like all nerve processes structurally related with the CBV wall, blind nerve endings are commonly surrounded by EF.

**Figure 29.**
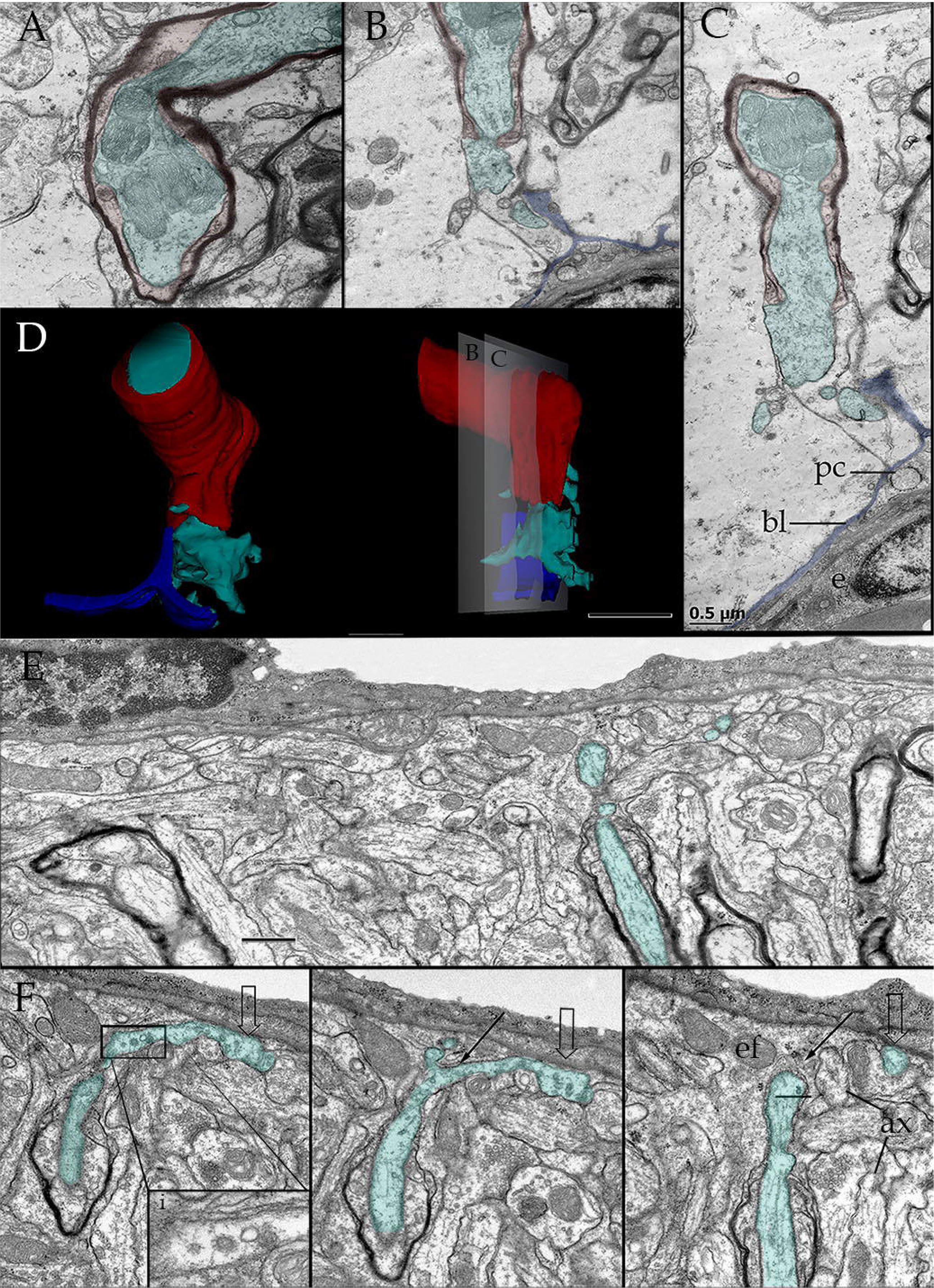
Series encompassing putative afferent or blind nerves in cerebellar cortex. A, B and C. Alternating micrographs through a myelinated nerve extending an bizarre outgrowth entirely engulfed by an end foot (ef). Notice that prior to losing the myelin envelope the axoplasm is filled by a cluster of mitochondria (m), that are, otherwise absent in the nude axoplasm (asterisk). bl = basal lamina; e = endothelium; pc = pericyte processes. D Reconstructions of the terminal axon that having lost the myelin envelope (red) expands (green) in that perivascular area adjacent to a longitudinal crest arising from the outer basal lamina (blue). E. Three terminal axons as assessed in a series encompassing them. A nude, narrow axoplasmic outgrowth (green) approaching the capillary wall of one of them (green). F. Subsequent series of that section shown in “E” defining the full distal axon trajectory to resolve (hollow arrow) next to the outer basal lamina surrounded by an end foot (ef). i. high magnification of the axoplasm containing coated vesicles. Notice the axoplasmic outgrowth of another nearby blind axon. (ax). Calibration bars: 0.5 µm.

The findings of this work extend previous observations revealing novel structures in the perivascular neuropil in several brain regions of diverse mammalian species that we propose are involved in mechanosensory functions. Addressing the possible source of the specialized synaptic terminals that innervate the LCs, we analyzed the cerebellar cortex. The above description suggest that in this brain region, glutamatergic neurons are a source of projections to the perivascular neuropil structures characterized herein. Hence, we searched for vesicular glutamate terminals by immunostaining on the cerebellum of transgenic animals expressing the GFP reporter under the gene control of the astrocyte marker GFAP (Figure 30). Confocal image stacks allowed 3D reconstructions that reveal the presence of glutamatergic terminals in close apposition to tunnels that we infer correspond to CBVs and enveloped by astrocytic cell extensions (Fig. 30). Some such terminals were located adjacent to nuclei that could corrrespond to pericytes and others seemed to be distant from these cells. Hence, these findings support the main notion of this work confirming the presence of what appears to be a tripartite synapse in the perivascular region.

**Figure 30.**
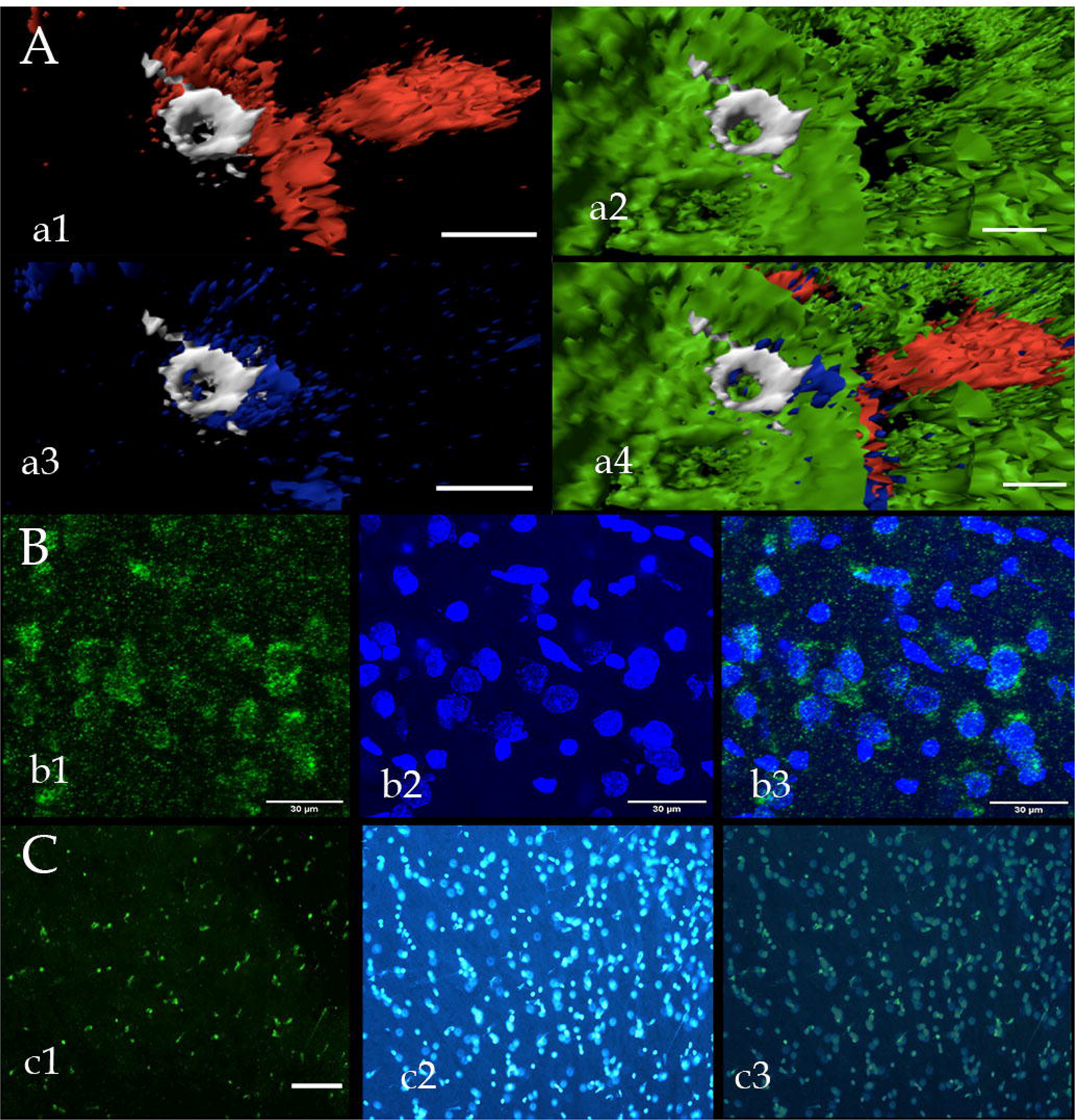
In situ hybridization and confocal, immunohistochemistry. A. Triple inmmunocytochemistry to CGRP (red) B4 isolectin (IB4)(white), vGlut2 (blue) and eGRP (green). a1 endotheliumm (white) CGRP, a2 eGRP and IB4, a3. IB4, a3. vGlut2 and IB4, a4 croop. B. In situ hyblidization to mRNA to Piezo 1 in layer V of the parietal lobe. b1 mRNA-Piezo1, b2 nuclei, b3 crop. C. In situ hyblidization to mRNA to Piezo 1 in the cerebellum deep granule cell layer. c1 mRNA Piezo1 probe. c2 nuclei. c3 croop.

Moreover, according to our findings, perivascular mechanosensory structures are involved in detecting changes in blood pressure in the CBVs. Recent discoveries have shed light in the age-long question of the molecular mechanisms that transduce mechanical stimuli to neural signals as part of diverse afferent systems. The proteins piezo1 and piezo2, ion channels whose activity is modulated by mechanical influences, have been found to be key players in this issue. Therefore, we analyzed the expression of piezo1 in the rat brain telencephalon by in situ hybridization. We observed the presence of piezo1 mRNA in the perinuclear region of cortical pyramidal (Fig. 30B) and thalamic (not shown) neurons. Given the fact that nearly the entire poblation of synapses containing small round vesicles arise from the later i.e., glutamatergic neurons (Bopp et al., 2017), it is highly plausible that the synaptic boutons and associated GCLs observed in tha BCU correspond to mechanosensory elements (Fig. 30B). Moreover, we observed expression signal in small clusters of piezo1 mRNA matching in location and dimensions cerebellar mossy fiber-rossetes (Fig. 30C). Of these structures, some had punctated signal that could correspond to mRNA in the axon terminals, while others revealed signal consistent with expression in the endothelial cells themselves. Hence, these findings are consistent with a role of Piezo 1 in mechanodetection in the brain and also reveal a higher complexity in the feed-back system that regulates microcapilar blood flow which warrants additional studies.

## Discussion

The fine structure and relevant immunohistochemical and molecular biological features of the brain capillary blood vessels and surrounding neuropil are presented. The main contributions of the present study are framed into a general description of the main structures conforming the neurovascular unit. The first relates to the general organization of the pericyte (PC) and its processes known to be a fundamental cell type for the genesis and blood flow regulation (Hirase et al, 2004) of blood vessels. Previously defined as the cell-type that encircles the capillary blood vessel (CBV) itself (Zimmermann, 1923), herein a novel pericyte (PC) process, or axonoid (AN) bridging heteronymous CBVs of uneven diameter was characterized. The discovery of a free PC lacking CBV and surrounded by neuropil and with a typical AN prompted the idea that it corresponds to a neuron, however, typical PC processes penetrate CBVs at either side. Also pertaining to the PC is another ubiquitous specialization, the asymmetric process (AP), consisting of a membranous structure associated with a blood flow impact sites (McDonald and Larue 1983). The AP is first described and fully characterized here. A modified tripartite synapse, (mTS) surrounded by EF (Reichenbach and Wolburg 2008) was found to be highly differentiated and crucial for the neural and glial interactions with the CBV. Thus, an earlier, masterful observation (Araque et al., 1999, Perea and Araque 2000) coupled with the observation that EF is the perivascular source of the third TS component (Cohen, et al., 1996, 1997, Reichenbach and Wolburg 2008, Varela-Echavarría et al., 2017). A previously unnoted difference is that perivascular synaptic terminals originate sets of tiny fibrils, termed here golf club-like extensions (GCLs). It is assumed that the complex GCLs link perivascular boutons with the outer basal lamina (oBL). More specifically, with a unique outgrowth of the latter termed here longitudinal crest (LC). The strategical localization of the EF in association with the synapse-GCL and LC supports a functional role. Indeed, it is likely that the well-known ability of peripheric astrocytic processes to control availability of synaptic glutamate, is shared by the EF. If so, Ca^++^ oscillations influenced by the blood flow may modulate the pericapillary synapse. 3D reconstructs showed that GCLs arising from series of perivascular synapses and are aligned longitudinally in the same order as parent boutons in the dendritic shaft (Fig. 16 and 19). It is suggested that the dendritic segment receiving series of consecutive TSs and their GCL along the LC and underlying PC furnish the so-called the neuro-glial capillary unit (NCU). With this background we were persuaded to search for the cell-types source of the unique pericapillary neuropil described here. Owing to the large number of short-axon neurons in the mammalian cerebral isocortex and thalamus we selected the cerebellar cortex, having a relatively small number of known neurons (Ramón y Cajal, 18989). Furthermore, the comprehensive description of the neuropil by Palay and Chan-Palay (1974) rendered feasible to accomplish the present ultrastructural correlates. Synaptic terminals of set of afferent fibers, i.e., mossy fiber-rosettes, granule cell perikaryon and its processes, and the interneuron axons exhibit varying degrees of apposition to the oBL. Like the NCU, cerebellar tributaries to the pericapillary domain were, invariably, encased by EF. The same applied for a small set of myelinated axons, resembling sensory fibers (Banks, 1986). The working hypothesis that the BCU represents the afferent link to the motor vascular neuron pool led us to search first for the putative neurotransmitter associated with TS-GCLs. Synaptic boutons with clear, rounded vesicles establishing asymmetrical contacts with dendrites is considered to be the structural signature of glutamatergic synaptic transmission. Hence immunoreactive sites (ir) to vGLUT 2 in the mouse cerebellum of the GFAP-GFP mouse with fluorescence in astrocytes (Nolte et al., 2001) were first performed. Furthermore, the expression of Piezo 1 was visualized by in situ hybridization. Focal perivascular and pyramidal cell Piezo 1 mRNA was found in thalamus and cerebral cortex. The finding that Piezo 1 mRNA is contained by cerebellar mossy fiber-rosettes which reperesent a set of intimate CBV interactions supports the direct involvement of mechanoreception in modulating the interneuron-projecting cell outcome. The role of the NCU in the broad context of cerebral blood flow and the plausible implications for Piezo 1 are discussed.

First to be commented upon are observations of the CBV-PC interactions revealing novel structural peculiarities. From the early thorough description of the PC of euplacental mammals (Zimmerman, 1923), it was known that this cell type organizes a flat, nearly continuous network encircling central and peripheral blood vessels. Although revising the vast current knowledge of the PC is beyond the scope of the present study, this cell type is fundamental in vascular genesis, metabolism, and vasomotor activity. Contiguity of the PC with arterial smooth muscle fibers (SM), another mural cell (Rouget, 1874), has led to the assumption that the PC subserves a vasomotor role, leading to a long-lasting debate (vide supra)(for reviews see Iadecola and Nedergaard, 2007, Winker et al, 2011). Interestingly, direct in vivo observations of CBVs are consistent with the existence of subsets of annular mural cells expressing SM antigens alternating with longitudinal cells lacking such markers (Hill et al., 2015). This dominant organization supports the idea that the PC provides a sensory network for exchange of information between neural and vascular compartments (Hill et al., 2015). The interlacing PC processes of an electrically excitable cell (Helbig et al., 1992) sealed by occasional gap- and numerous tight junctions, provides a potential physiological link between PCs. Notably, interlacing PC processes protruding underneath the oBL-LC and united by gap- and tight junctions suggests a joint functional interaction (vide infra) as well. However, until the precise physiological involvement of the PC-CBV is known, implications for putative PC connectivity will remain an open question. In the same context falls the tangential route defined by the perforating ANs and PCs described here. Resembling an axon, the AN also leaves the parent PC-CBV and pierce the neighboring neuropil. Unlike the axon, however, the AN anchors on a PC encircling a CBV of dissimilar diameter. Our observations add new elements to the description of a perforating PC process in the human cerebral cortex (Shapson-Coe et al, 2021). Namely, the identification of the AN as a distinct PC process, the asymmetry of CBVs linked by perforating ANs, and the PC with an outstanding location in the neuropil proper. The AP, a distinct membranous component was added to the PC (Figure 10). Following its initial description with the Golgi technique, the electron microscope added several intrinsic and associated structures (Fig. 12) all of which appear to be unified by a common functional, possibly a sensory role. This provides a substratum to the long-standing dilemma on the reversible vasomotor activity elicited by functionally recruited areas of the brain, the so-called BOLD effect (Tsvetanov et al., 2020). Validity of these correlates ultimately rely on the ability of the NCU to decode the subtle mechanical or chemical stimuli generated by the blood flow, an open question amenable to future physiological assessment.

The NCU is a highly specialized differentiation of the CBV-neuropil. It occurs in certain segments of the CBV characterized by EF covering discernible with the light microscope (Fig. 7B). Moreover, in our specimens, obvious gaps denuded of EF were commonly observed as well (Figs.5C and 15A) which contrasts with the complete EF covering of blood vessels with diameters larger than 15 microns (Mathiisen, et al., 2010). A first exploration revealed a high density of NCU in isocortex and thalamus. Fundamental evidence to support the sensory involvement of the NCU in these brain regions is a high incidence of associated mTSs. Indeed, the starting point of the present work was the observation that sets of TSs were consistently engulfed by EF (Varela-Echavarría et al., 2017). The involvement of pericapillary EF synaptic interactions (Cohen et al., 1996, Cohen et al 1997) and of the EF itself responding to blood flow (Paulson et al., Neuman et al, 1984) and influencing both CBV diameter and the ensuing vasomotor response (Zonta et al., 2003) biased our search for interactions between synapse-EF and the CBV proper. Thus, the presence of tiny tubular structures trapped by the EF cytoplasm (Fig. 18C-F) prompted the use of serial sections. It was soon found that the enigmatic fibrils piercing the EF correspond to extensions of the synaptic bouton organizing the TS. The ubiquity of the now termed GCLs along with their alignment to the LC and to the opposing alpha and beta AP processes with a distinct ground substance (Schwartz and DeSimone 2008, Vega et al., 2009) is reminiscent the row of hairy receptor cells aligned with the tectorial membrane or with the macula saculi of primary hearing and balance organs (Kahle et al., 1978), all mechanical transducing structures. Hence, the GCLs necessitate transducing elements to yield to mechanical energy EF membrane Ca^++^ oscillations (Neuman et al 1984, Paulson and Neuman 1987, Windship et al, 2007, Deitmer et al., 2009) or receptor potentials (RPs), and neuron retrieval (Shmuel et al 2002, 2006, Tsvetanov, 2020, Vaucher et al., 2000, Anenberg et al,, 2015). An important clue in this regard was provided by the vesicle content of both perivascular synaptic bouton-GCL typical of glutamatergic, excitatory synapses (Peters et al., 1974, Uchizono 1975). Indeed the EF surrounding a series of axo-dendritic boutons originating a likewise ordered set of GCLs next to the oBL-LC suggests p*er se* a sensory role. A fair interpretation of this is that mechanical stimuli that modify the EF Ca^++^ oscillations modulate synaptic transmission (Schipke and Kettenmamm, 2004) and may also impact the GCL-synaptic output to its dendritic tributary or to the parent axon. Although the definition of the chemical nature of the synaptic GCL (Sy-GCL) is still in process, the vesicle content coupled with our immunohistochemical observations support its glutamatergic nature.

Relevant to present correlates would be the existence of baroreceptor ion-channels within the perivascular domain. This prompted us to analyze the expression of the Piezo 1 mechano-sensitive channel gene by in situ hybridization. The light microscopic evidence supports the existence of mRNA throughout the capillary wall (not shown), that is observable in large glutamatergic synaptic complexes such as the mossy fiber. This indirect evidence, coupled with the ubiquitous distribution of Piezo 1 and 2 mRNA in vascular endothelium (Li et al., 2014) outside the brain lead to the possibility that the brain CBV endothelium is responsible for the mechanical transduction of the NCU or that Piezo 1 mRNA is present in the Sy-GCL. The best argument in favor of the latter is the astonishing Piezo 1 expression in pyramidal (Fig. 30) and thalamic neurons, the main neuronal source of cortical synaptic glutamate. Therefore, being pyramidal cells the most likely source of the Sy-GCL, it is reasonable that mechanical disturbances impacting the CVB wall are transmitted by the EF low viscosity (Lu et al., 2006) eliciting Ca^++^ signaling which would recruit would recruit the aligned Sy-GCLs ultimately influencing dendritic processes of the neuron motor pool (Shmuel 2002, 2006, Hill et al., 2015). Aside of the present postulate requiring further immunocytological- and physiological assessment, another basic dilemma arises regarding the neuronal identity of axon and dendritic tributaries to the dendrite-Sy-GCL.

While the foregoing remarks pertain to the putative sensory role of the NCU, a possible afferent-efferent dual role should also be considered as observed in most peripheral sensory organs (Banks 1986, Banks 2015; Bewick and Banks 2015). Indeed, the early identification of synaptic vesicles in primary afferent axons constituted a puzzling observation (Kruger 1988). Electron microscopic studies with vesicle tracing techniques, however, have demonstrated that a set of synaptic vesicles in sensory nerve terminals actually correspond to recycling elements. Furthermore, ulterior cytochemical work provided direct evidence for vGlut2 release enhancing the receptor output (Bewick and Banks 2015). The omnipresence of coated vesicles associated with positivity to vGut2 support their involvement in tune-up the receptor proper. However, subsequent observations (Bopp et al., 2017) in the sensory and motor cortex indicate that synaptic structure is comparable, irrespective of the presence or absence of vGlut 2-immunoreactivity. Thus, the possible presence of vGlut2 immunoreactive sites associated with the NCU supports a reciprocal between the GCL-synaptic bouton with the capillary wall proper. Therefore, we performed combined immunohistochemistry to vGlut2 and to CGRP, a landmark molecule for sensory receptor axon identification (Silverman and Kruger, 1989) and blood flow modulation (Brain and Grant, 2004). Our studies revealed regions in which vGlut2 and GCRP are present in sites corresponding to the NCU. Hence, the presumptive mechanotransduction ascribed to the NCU may be subject to a vGlut2-mediated reciprocal modulation at the CGRP immunoreactive sites delivered by ascending i.e., afferent tracts, presumptive source of GCL-synapse (vide infra).

Although the presence of synaptic terminals to dendrites within the CBV-neuropil intersection is per se relevant, it is perhaps equally important to define their parent neurons. To this end, we selected the cerebellar cortex as its main afferences and neuron types (Ramón y Cajal 1904) as well as the ensuing neuropil have been thoroughly studied (Palay and Chan-Palay 1974, Hamory and Somogyi 1983, Sotelo 2008) affording base line knowledge to perform electron microscopic interpretation. The Golgi technique utilized first, revealed axonal contributions of both arriving and intrinsic cortical cerebellar connections. In fact, the CBV contribution should be added to Caja’s basic circuitry. For instance, in comparing the axonal paths traced by the granule cell (GC) to the molecular layer, we found a unique collateralization of the GC to the CBV, although a notable somatic-dendritc contribution was also evident. While the wealth of information provided by the Golgi-technique could not be correlated with our electron microscopic observations, a necessary first step to the organization of the cerebellar perivascular neuropil is offered. Thus, ascending fiber systems defined by the mossy-, climbing-(Fig. 10A), and parallel fibers display obvious, chiefly in passage contribution to the pericapillary neuropil. To be sure, all of them represent the glutamatergic source of axon terminals to the interneuron-projecting cell chain. Therefore, the possibility arises that the volleys of nerve impulses arriving to the cerebellar cortex modulated by the EF-CBV interaction so that the red blood cell column impacts (Noguchi and Gommpper 2005 Moore and Cao 2008) might influence the eventual synaptic outputs of parent axons providing pericapillary boutons? The possibility was assessed by searching for Piezo 1 and Piezo 2 mechanosensitive channels, whose known ganglion cell expression is necessary to elicit the homeostatic baroreflex (Zeng, et al., 2018). Morphological insight for Piezo 2 involvement in brain mechanic sensitivity was the immunohistochemical demonstration of Piezo 2 immunoreactivity in principal cells (Wang and Hamil 2020) and deep cerebellar nuclei (Wang and Hamil 2021). The existence of endothelial mechano-sensitive Piezo 1 channels in peripheral blood vessels (Li et al., 2014) and Piezo 2 in the ganglion cell-mechanoreceptor unit (Woo, et al 2015), and the frequent CVB interaction with mossy fiber-rosettes led us to further investigate a possible Piezo 1 baroreceptor channel associated to them. Hence, we cloned the Piezo 1 gene and provide molecular evidence (i.e., in situ hybrization) for expression in isocortex pyramidal cells and in small aggregates matching the cerebellar mossy fiber-rosette in both size and distribution. Overall that the ability of capillary blood flow to elicit a reciprocal response in the ascending MF-R would influence the activity of the short-axon and Purkinje principal cell.

### Concluding remark

The existence of cytological and molecular substrates transducing endogenous mechanical stimuli delivered by the blood flow is highly relevant to account for fundamental aspects of the functioning of the nervous system. Hence, the putative hemodynamics sensing structures described herein provide an essential mechanistic element for any task requiring neuronal circuits decoding blood flow requirements or rhythmical events such as pulsating blood flow. Experimental testing of this postulate in future studies is feasible with electrophysiological, molecular, pharmacological or behavioral strategies.

## Acknowledgements

The 3D image processing was performed at the Laboratorio Nacional de Visualización Científica Avanzada (LAVIS) with the support of Alejandro de León Cuevas, Alejandro Ávalos Fernández and Luis A. Aguilar Bautista. Technical support was also received from Adriana González Gallardo, Michael Jeziorsk, Ma. Antonieta Carbajo, Loudes Palma, Martín García Servín.

## Notes

### Competing Interest Statement

The authors have declared no competing interest.

### Summary of Updates

Figures were moddified according with the bioRxiv policies

